# Advanced human iPSC-based modelling of *LMNA*-related congenital muscular dystrophy enables development of targeted genetic therapies for muscle laminopathies

**DOI:** 10.1101/2025.06.22.660928

**Authors:** Daniel Moore, Heather Steele-Stallard, Luca Pinton, Valentina Maria Lionello, Lucia Rossi, Artadokht Aghaeipour, Salma Jalal, Cherry Tsz Yan Wong, Angela Clara-Hwang, Gisèle Bonne, Peter S. Zammit, Francesco Saverio Tedesco

## Abstract

*LMNA*-related congenital muscular dystrophy (L-CMD) is amongst the most severe forms of laminopathies, which are diseases caused by pathogenic variants in the *LMNA* gene. *LMNA* encodes the proteins Lamin A and C, which assemble with Lamin B1 and B2 to form the nuclear lamina: a meshwork providing structural stability to the nucleus that also regulates chromatin organisation and gene expression. Research into L-CMD mechanisms and therapies is hindered by lack of humanised, tissue-specific models that accurately recapitulate disease phenotypes. We previously reported that *LMNA*-mutant induced pluripotent stem cell (iPSC)-derived skeletal muscle cells have nuclear shape abnormalities and Lamin A/C protein mislocalisation. Here, we expand the selection of L-CMD patient-derived iPSCs and validate disease-associated readouts using a transgene-free based protocol which more accurately mimics skeletal myogenesis. Results showed no overt defects in developmental myogenesis, but recapitulation of pathological nuclear shape abnormalities in 2D and 3D cultures, nuclear envelope protein mislocalisation and transcriptomic alterations across multiple pathogenic *LMNA* variants. We then utilised this platform to assess *LMNA* gene editing strategies. CRISPR-based exon removal generated stable RNA and protein Lamin A/C species, without significant normalisation of nuclear morphological phenotypes or transcriptomic profile. Conversely, precise editing of the same mutation showed complete reversal of disease-associated nuclear morphometrics, alongside normalisation of the pro-inflammatory transcriptomic signature. Our data provide the foundation for a humanised *in vitro* disease and therapy modelling platform for this complex and severe muscle disorder.

**GRAPHICAL ABSTRACT:** 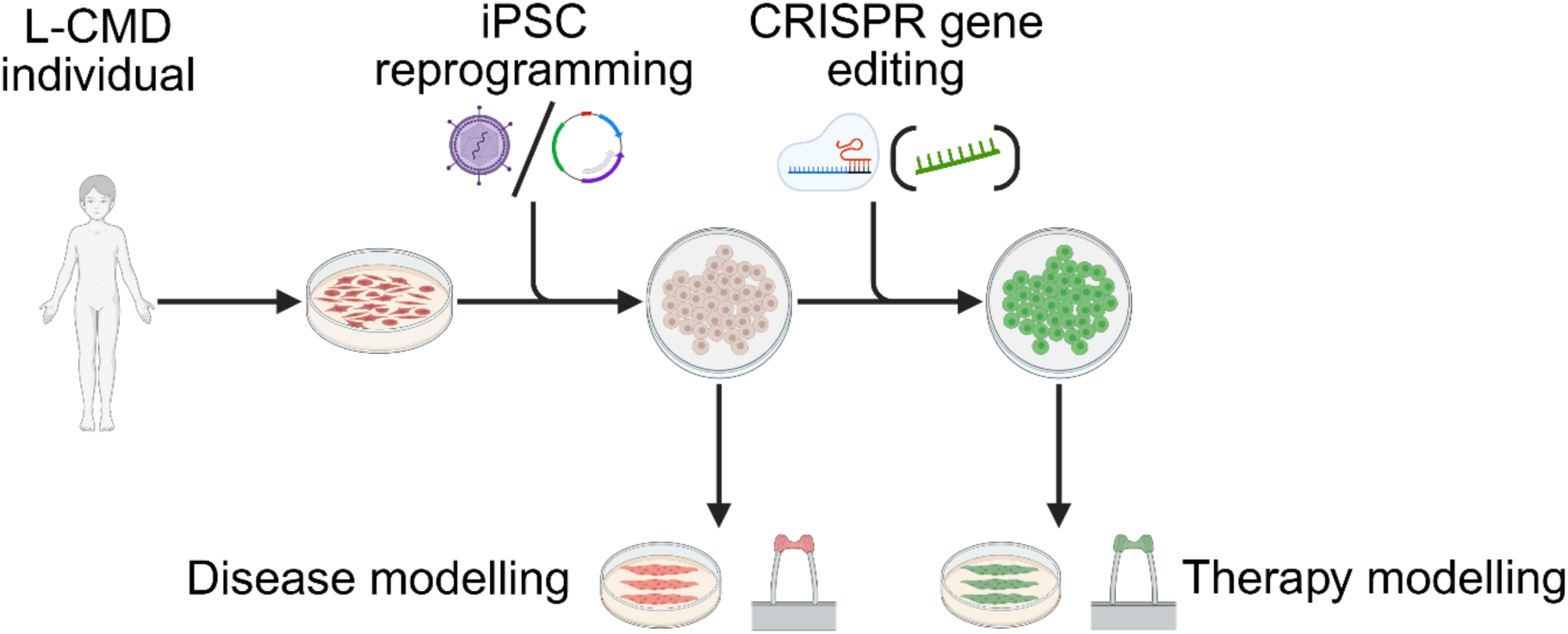

**HIGHLIGHTS:**

- *LMNA*-mutant iPSCs undergo efficient skeletal myogenesis upon transgene-free, small molecule-based lineage-directed differentiation
- L-CMD iPSCs recapitulate hallmark disease-associated nuclear phenotypes and show a pro-inflammatory transcriptional profile
- Disease modelling platforms based on iPSC-derived skeletal muscle cells enable comparative testing of gene editing strategies
- CRISPR-edited L-CMD iPSC-derived myogenic cells show amelioration of disease-associated readouts

## INTRODUCTION

Laminopathies are a group of genetic diseases caused primarily by mutations in the *LMNA* gene. *LMNA* encodes Lamin A/C, which assemble with Lamin B1 and B2 to form the nuclear lamina: a meshwork providing structural stability to the nucleus^1^. At least 16 diseases are caused by mutations in the *LMNA* gene, amongst which the most common and severe are those affecting striated muscle tissues. In particular, *LMNA*-related congenital muscular dystrophy (L-CMD) is a severe paediatric form of cardiac and skeletal striated muscle laminopathy, clinically characterised by early onset muscle weakness and wasting and dropped head syndrome. Sudden death in the first decade of life is common due to cardio-respiratory failure^2,3^. The precise pathological mechanisms of laminopathies remain elusive and likely varies by mutation and individual.

Two non-mutually exclusive hypotheses have emerged as primary candidate mechanisms for pathogenesis. The mechanical stress hypothesis focuses on the breakdown of the mechanoresponsive properties of the nuclear lamina when mutated. A key phenotype of laminopathies is presence of abnormally shaped nuclei, which has been modelled in *in vitro* and *in vivo* studies^4–12^. Impaired mechanoresponse can lead to mispositioning of nuclei within muscle tissues, potentially perturbing myonuclear domain positioning^7,13^. Furthermore, many *LMNA*-mutations increase susceptibility of nuclei to rupture, the functional consequences of which can include loss of genetic and nucleoplasmic material, activation of inflammatory response pathways and cell death^14–17^. Ultimately, these mechanisms may be particularly relevant in mechanically active tissues such as striated muscle, which is the most affected tissue in laminopathies.

The gene regulation hypothesis states that pathological changes in transcription occur due to altered chromatin organisation and contact with mutant Lamin A/C. Studies across various model systems and laminopathy subtypes have revealed alterations to Lamin A/C-chromatin interactions at lamina-associated domains, gene enhancers, and transcription start sites, as well as altered interactions with the epigenetic repressive polycomb group proteins^18–23^. Two studies have examined transcriptional dysregulation caused by *LMNA* mutations in skeletal muscle cells. The first, revealed alterations to signalling pathways such as MAPK, Akt/mTOR, AMPK and proteostasis, via RNA-sequencing of biopsied muscle tissue from a patient carrying the Lamin A/C p.G449V mutation^24^. More recently, a study identified disruption of metabolism, extracellular matrix and fibrosis, de-repression of alternative fate genes and disrupted splicing pathways as integrative mechanisms of pathogenesis in primary myoblast lines from patients with Emery-Dreifuss muscular dystrophy (EDMD), caused by mutations in *LMNA*, *TMEM214*, *PLPP7*, *SUN1*, *SYNE1*, *EMD* and *FHL1*^25^. However, the impact of variants causative of L-CMD (the most severe striated muscle laminopathy) on the transcriptome remains unexplored, possibly due to the invasive sampling required in such a severe, early-onset condition.

While these studies highlight general pathways and mechanisms of laminopathy pathogenesis, the lack of effective, humanised models of L-CMD prevents examination of tissue and disease-specific pathomechanisms. A concern when using non-human models of laminopathies is the inability of these models to recapitulate the precise pathogenetic events leading to disease. For example, murine models typically require homozygous *LMNA* mutations to elicit a phenotype, whereas the majority of human mutations are heterozygous, with a dominant-negative disease mechanism^26^. Therefore, humanised model systems using primary, immortalised, or iPSC-derived myogenic cells are being used to investigate tissue and disease-specific pathomechanisms of laminopathies. Despite this, research into L-CMD using humanised model systems remains limited. Primary and immortalised human myoblasts have been used to study mechanotransduction defects and nuclear shape changes in miniaturised engineered muscle models^4,27–30^. However, given that L-CMD is a rare disease and no longer requires invasive muscle biopsy for diagnosis, patient-specific cell lines are often limited in number and difficult to access for the wider research community. To overcome this limitation, we have previously harnessed the controllable proliferative and differentiative capacity of iPSCs to study laminopathies in a humanised, tissue-specific manner in monolayer and 3D engineered muscle tissues, enabling modelling of nuclear shape abnormalities and providing evidence for the potential of iPSCs in laminopathy research^9,12^. Nonetheless, the full potential of *LMNA*-mutant iPSCs to answer key questions in laminopathies, such as genotype-phenotype correlations, impact on human skeletal myogenesis, transcriptional alterations and capacity to predict efficacy of novel therapeutic strategies, remains unexploited.

Here, we expand our iPSC-based modelling platform to include a novel L-CMD line and examine the impact of different *LMNA* mutations on myogenesis, model nuclear morphological abnormalities and protein mislocalisation phenotypes and analyse transcriptomic alterations in *LMNA*-mutant iPSC skeletal myogenic derivatives in 2D and 3D culture systems. Furthermore, we use this iPSC-based platform to examine multiple CRISPR-Cas9-based therapeutic approaches for the amelioration of disease-associated phenotypes in L-CMD.

## RESULTS

### Human iPSCs harbouring pathogenic *LMNA* variants undergo efficient skeletal myogenesis upon transgene-free, small molecule-based differentiation

We previously reported that cellular features of skeletal muscle laminopathies can be modelled using human iPSCs carrying pathogenic *LMNA* variants differentiated into skeletal myogenic cells by overexpressing the myogenic regulator MYOD^9^. However, other studies have demonstrated impaired skeletal myogenic differentiation in *LMNA*-mutant or -null cells^5,31–33^. Therefore, to determine whether pathogenic *LMNA* variants interfere with human developmental skeletal myogenesis *in vitro*, we selected two readily available, fully characterised iPSC lines – *LMNA*^K32del^ (hPSCreg: CMDi003-A; K32del) and *LMNA*^R249W^ (hPSCreg: CMDi004-A; R249W) – alongside a novel *LMNA*^L302P^ iPSC line generated for this study (L302P; characterisation in Figure S1) and differentiated them into skeletal myogenic progenitors and myotubes using a transgene-free, small molecule-based protocol, modelling developmental myogenesis^34^. These 3 iPSC lines were associated with the severe L-CMD phenotype and included different types of pathogenic variants (i.e. R249W and L302P are missense, while K32del contains a deletion), frequency (relatively common for *LMNA*^R249W^ vs. ultra rare with *LMNA*^L302P^) and regions of the gene/protein affected (K32del affecting exon 1, R249W exon 4 and L302P exon 5)). Two healthy donor iPSC lines were used as controls.

We first confirmed the presence of the pathogenic *LMNA* variants (K32del, R249W and L302P; Figure 1A) and pluripotency-associated factors through immunolabelling for SOX2, NANOG and OCT4 (Figure 1B, C). Lamin A/C is known to be absent in pluripotent cells and then progressively expressed in most somatic tissues as differentiation proceeds^35^. As expected, undifferentiated iPSC colonies were negative for Lamin A/C (Figure 1B).

**Figure 1.**
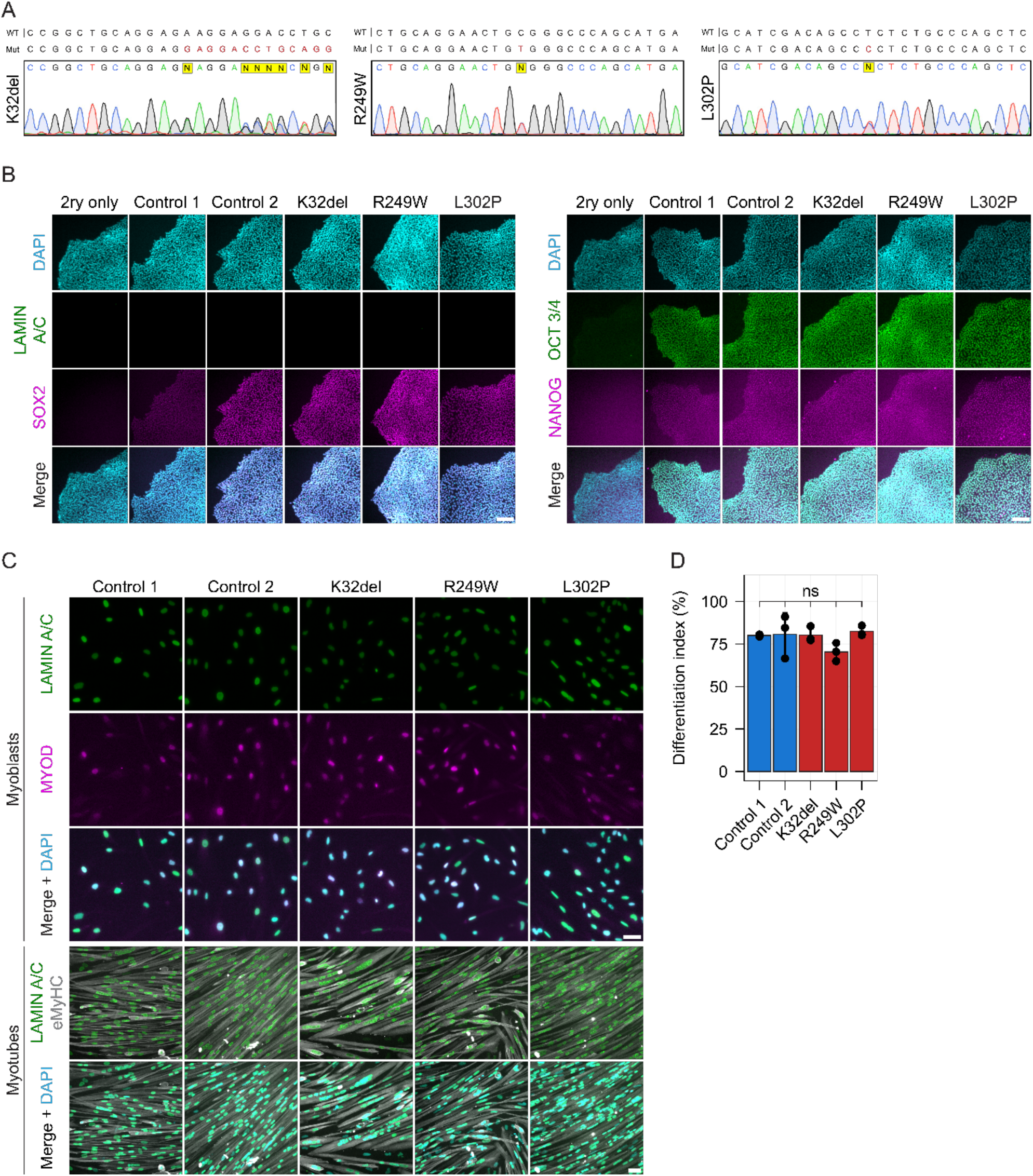
L-CMD iPSCs undergo efficient transgene-free, small molecule-based skeletal myogenesis *in vitro*. **(A)** Sanger sequencing electropherograms confirming the presence of heterozygous *LMNA* pathogenic variants in the cell lines used. **(B)** Immunostaining of NANOG and OCT3/4, SOX2 and Lamin A/C in iPSCs. Nuclei counterstained with DAPI. Scale bar 100µm. **(C)** Immunostaining of Lamin A/C and MYOD in control and *LMNA*-mutant myoblasts and of Lamin A/C and embryonic myosin heavy chain (eMyHC) in control and *LMNA*-mutant myotubes. Counterstained with DAPI. Scale bar 50 µm. **(D)** Differentiation index of control and *LMNA*-mutant myotubes. One-way ANOVA with Tukey’s post-hoc tests for comparisons to controls. Three independent passages were analysed per cell line (N = 3). 1311 – 3562 nuclei were analysed per cell line, per repeat. ns p>0.05. Dots show mean value of each replicate. Whiskers show standard deviation.

To confirm commitment to the myogenic lineage, we immunolabelled for the master transcriptional regulator of muscle differentiation MYOD. MYOD and Lamin A/C were detected in all cell lines, indicating successful myogenic commitment (Figure 1C). We then induced terminal skeletal myogenic differentiation by allowing cells to reach confluency, followed by serum starvation. Cells fused over the course of 2-4 days to form multinucleated, myosin-positive myotubes with Lamin A/C-positive myonuclei (Figure 1C). Interestingly, we observed that non-differentiated cells often expressed visibly less Lamin A/C as indicated by decreased fluorescent signal compared to myonuclei. To examine whether a differentiation defect was present in the *LMNA*-mutant lines, we analysed the differentiation index. Results showed no significant myogenic differentiation defect in any of the *LMNA* mutant lines versus controls (Figure 1D; mean ±SD: Controls 1 and 2 80.07 ± 0.60% and 80.68 ± 12.69% respectively, K32del, R249W and L302P 80.27 ± 4.53%, 70.35 ± 5.36% and 82.32 ± 3.09% respectively). These data show that dominant-negative *LMNA* variants do not impair exit from pluripotency and differentiation of human iPSCs into the skeletal muscle lineage *in vitro*.

### *LMNA*-mutant, iPSC-derived skeletal myotubes have abnormal nuclear morphology, nuclear envelope protein localisation and nucleoplasmic reticulum

Having efficiently generated *LMNA*-mutant skeletal myotubes from multiple L-CMD iPSCs, we then assessed presence of hallmark laminopathy nuclear shape abnormalities^9^. We observed a range of myonuclear dysmorphism in *LMNA*-mutant populations that we broadly categorised as “Normal”, “Deformed”, “Blebbed”, “Elongated” or “String” (Figure 2A; Figure S2A). To examine whether these categories were present in significantly larger numbers in *LMNA*-mutant cell lines versus controls, we quantified the proportion of each of these categories in a blinded manner (Figure 2B). Each *LMNA*-mutant line had significantly reduced percentages of “Normal” myonuclei versus controls (mean ±SD: Control 1 93.87 ± 3.32%; Control 2 95.11 ± 2.80; K32del 55.09 ± 4.75%; R249W 48.65 ± 5.45%; L302P 68.15 ± 3.21%). Overall, “Deformed” was the most common dysmorphic classification in *LMNA*- mutant myonuclei (mean ±SD: Control 1 5.32 ± 2.58%; Control 2 4.27 ± 2.26%; K32del 20.84 ± 4.09%; R249W: 25.19 ± 1.77%; L302P: 20.44 ± 1.46%). Blebbed was the next most frequent classification in K32del (mean ±SD: Control 1 0.80 ± 0.80%; Control 2 0.43 ±0.38%; K32del 18.47 ± 11.02%; R249W 9.42 ± 1.42%; L302P 4.93 ± 1.04%), while elongated myonuclei were the second most common dysmorphism in R249W and L302P (mean ±SD Control 1 0.13 ± 0.23%; Control 2 0.00 ± 0.00%; K32del 11.68 ± 2.31%; R249W 21.07 ± 6.41%; L302P 6.39 ± 4.49%). String myonuclei were the most severe and least common dysmorphism in all *LMNA*-mutant lines and the only one not observed in both Controls (mean ±SD: Control 1 0.00%; Control 2 0.00%; K32del 2.51 ± 1.55%; R249W 3.63 ± 1.29%; L302P 1.22 ± 0.64%).

**Figure 2.**
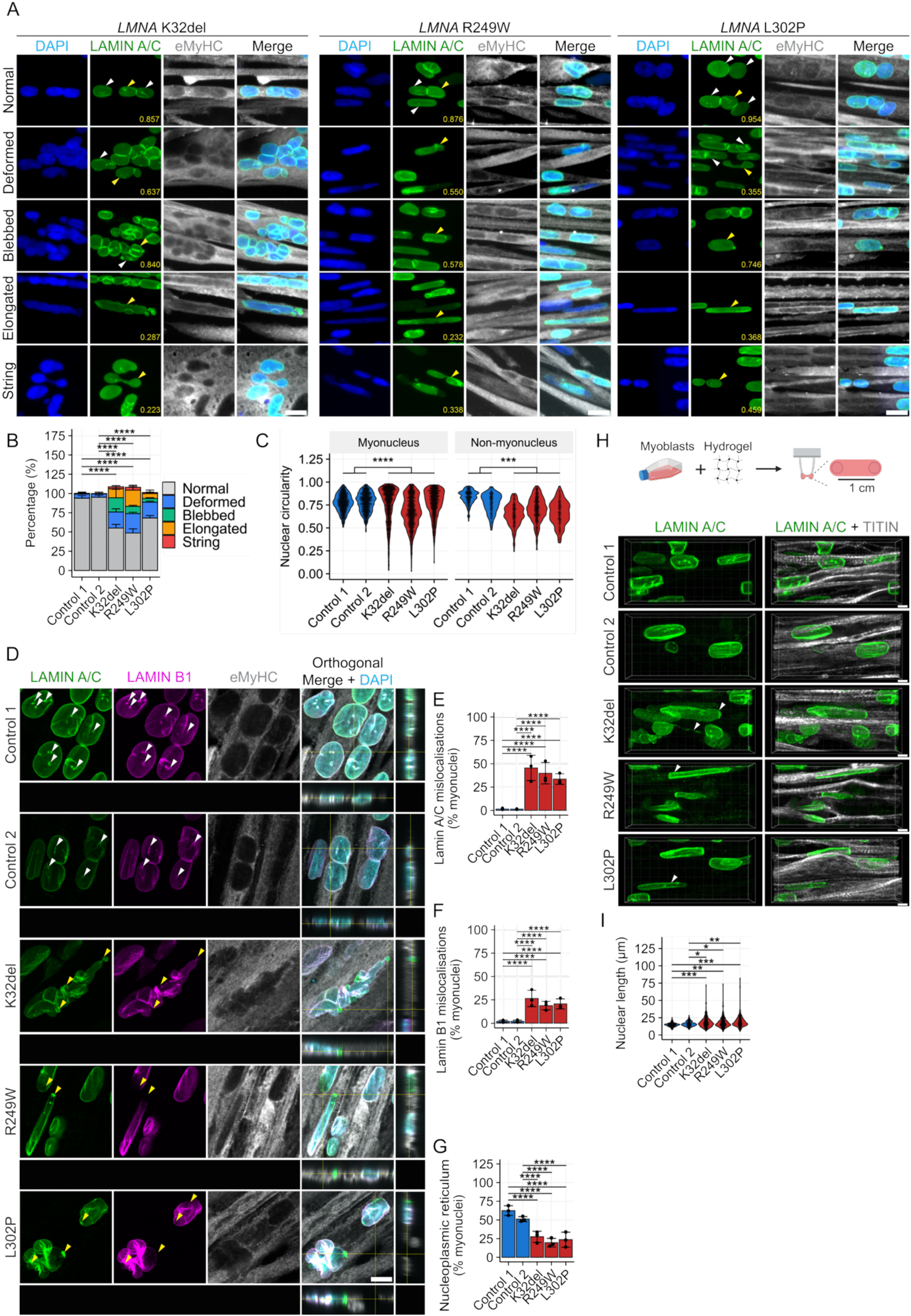
L-CMD iPSC-derived myonuclei show pathological laminopathy phenotypes in monolayer cultures and 3D engineered muscles. **(A)** Immunostaining of Lamin A/C and embryonic myosin heavy chain (eMyHC) showing representative images of five nuclear morphology classifications are displayed: Normal, Deformed, Blebbed, Elongated, and String. Arrows indicate nuclei belonging to shape classification. Yellow arrows indicate nucleus associated with nuclear circularity measurement in lower right corner. Scale bar 20 µm. **(B)** Proportion of nuclear shape classifications in myotube nuclei. Nuclei with more than one type of abnormality (e.g. elongated and deformed) were scored twice meaning totals can exceed 100%. Nuclei were considered elongated if they had a major axis length >30 µm. Number of normal shaped nuclei were compared between *LMNA*-mutants and controls for fractional logistic regression with Tukey’s correction for pairwise comparisons. Three independent passages were analysed per cell line (N=3). 96-291 myotube nuclei were analysed per cell line, per repeat. **** p<0.0001. Data presented as stacked bar chart. Whiskers show standard deviation. **(C)** Graphs showing quantification of nuclear circularity. Beta regression with Bonferroni correction for pairwise comparisons. Data presented as scatter plots showing measurements of all nuclei across repeats with violin plots showing distribution of measurements. Three independent passages were analysed per cell line (N=3). 205-622 myotube nuclei and 50-174 non-myotube nuclei were analysed per cell line, per repeat. *** p<0.001; **** p<0.0001. **(D)** Representative immunostain of Lamin A/C, Lamin B1 and eMyHC. Counterstained with DAPI. Z-stack images of control and *LMNA*-mutant nuclei are displayed as a single Z-slice, with orthogonal view. White arrowheads indicate nucleoplasmic reticulum, yellow arrows indicate Lamin A/C and Lamin B1 mislocalisations. Scale bar 20 µm. **(E, F)** Quantification of (E) Lamin A/C mislocalisation and (F) Lamin B1 mislocalisation. Fractional logistic regression with Tukey’s post hoc test for pairwise comparisons. Three independent passages were analysed per cell line (N = 3). A total of 119 – 347 nuclei were analysed per cell line, per repeat. **** p<0.0001. Dots represent the mean value of one replicate. Whiskers indicate the standard deviation. **(G)** Quantification of nucleoplasmic reticulum positive nuclei in control and *LMNA*-mutant myotube nuclei. Fractional logistic regression with Tukey’s post hoc test for pairwise comparisons. Three independent passages were analysed per cell line (N = 3). 107 – 246 nuclei were analysed per cell line, per repeat. **** p<0.0001. Dots show mean value of one replicate. Whiskers indicate standard deviation. **(H)** Schematic diagram showing generation of 3D engineered muscles and immunostain for Lamin A/C and Titin in 3D engineered muscles. Arrows indicate elongated myonuclei. Scale bar 5 µm. **(I)** Graph showing quantification of myonuclear length in 3D engineered muscles. One-way ANOVA with Tukey’s correction for pairwise comparisons on log-normalised data to achieve normal distribution. Data presented as scatter plots showing measurements of all nuclei across repeats with violin plots showing distribution of measurements. Three independent passages were analysed per cell line (N=3). 44 - 302 nuclei were analysed per cell line, per repeat. * p<0.05; ** p<0.01; *** p<0.001.

To complement this analysis, we quantified additional nuclear morphometric parameters. We assessed nuclear circularity ratio (NCR)^9^ and *LMNA*-mutant cell lines had significantly less circular nuclei than healthy controls (Figure 2C; mean ±SD: control myonuclei 0.779 ± 0.016; control non-myonuclei 0.805 ± 0.042; *LMNA*-mutant myonuclei 0.731 ± 0.053; *LMNA*-mutant non-myonuclei 0.670 ± 0.047). In *LMNA*-mutant cell lines, a population of severely dysmorphic myonuclei and non-myonuclei were observed, corresponding to NCR scores of <0.5, which were typically not present in control cells. Nuclear perimeter and nuclear area were also measured, revealing significantly increased myonuclear and non-myonuclear perimeter in *LMNA*-mutant lines (Figure S2B; mean ±SD: control myonuclei 47.92 ± 2.67 µm; control non-myonuclei 59.91 ± 4.30 µm; *LMNA*-mutant myonuclei 53.77 ± 3.24 µm; *LMNA*-mutant non-myonuclei 71.69 ± 5.45 µm). We did not observe an increase in nuclear area in either myonuclei or non-myonuclei between *LMNA*- mutants and controls (mean ±SD: control myonuclei 143.13 ± 16.99 µm^2^; control non-myonuclei 233.21 ± 25.83 µm^2^; *LMNA*-mutant myonuclei 159.06 ± 14.97 µm^2^; *LMNA*-mutant non-myonuclei 270.89 ± 28.96 µm^2^; Figure S2C).

Cellular phenotypes of laminopathies also include mislocalisation of nuclear lamins, with striated muscle laminopathy-causing mutations having a particular propensity to form Lamin protein aggregates^36^. Investigating Lamin A/C and Lamin B1 mislocalisation showed each *LMNA*-mutant cell line exhibited mislocalisation of both Lamin A/C and B1, which were typically characterised by aggregations of protein or localised depleted immunosignal (Figure 2D). Lamin A/C mislocalisation was minimal in the control lines (mean ±SD: Control 1 1.57 ± 0.81%; Control 2 1.46 ± 0.38%), whereas all *LMNA*-mutant lines showed significantly elevated levels (mean ±SD: K32del 45.51 ± 13.51%; R249W 39.81 ± 11.40%; L302P 33.57 ± 5.65%; Figure 2E). Similarly, Lamin B1 mislocalisation levels were low in control lines (mean ±SD: Control 1 2.06 ± 1.19 %; Control 2 2.04 ± 1.30 %) and were significantly increased in *LMNA*-mutant lines (K32del 26.48 ± 8.78 %; R249W 18.85 ± 4.51 %; L302P 20.70 ± 5.41 %; Figure 2F).

We then examined the nucleoplasmic reticulum in myonuclei of the same cultures. These structures are invaginations of the nuclear envelope which have been proposed to play roles in calcium signalling, structural support and transport of small molecules across the nuclear envelope^37,38^. These structures are lined by the nuclear lamina and as such, we hypothesized that a destabilisation of nuclear lamin proteins may lead to the formation of less nucleoplasmic reticulum. We observed significantly reduced numbers of myonuclei with nucleoplasmic reticula in all *LMNA*-mutant lines assessed versus both controls (mean ±SD: 62.46 ± 6.52 % and 51.07 ± 3.44 % in Control 1 and 2 respectively; 27.47 ± 7.47 %, 19.55 ± 5.73 % and 23.57 ± 10.04 % in K32del, R249W and L302P, respectively; Figure 2G).

We previously showed that modelling laminopathies in 3D engineered muscle tissues exacerbates myonuclear dysmorphic phenotypes in *LMNA*-mutant cells^9^. We therefore generated 3D engineered muscle tissues using transgene-free iPSC-derived myoblasts to validate this observation and study nuclear morphology in a three dimensional environment closer to native tissue architecture (Figure 2H; Figure S2D). Comparison of *LMNA*-mutant engineered muscles to controls revealed a myonuclear elongation phenotype (mean ±SD: Control 1 14.69 ± 0.12 µm; Control 2 15.75 ± 0.44 µm; K32del 17.81 ± 0.19 µm; R249W 17.59 ± 0.60; L302P 18.85 ± 0.86 µm; Figure 2I). Together, our results validate myonuclear shape and protein mislocalisation as robust phenotypes across iPSC myogenic differentiation protocols, supporting the use of *LMNA*-mutant iPSC derivatives in disease modelling for laminopathies.

### Transcriptomic analysis of *LMNA*-mutant skeletal myogenic iPSC derivatives reveals a pro-inflammatory profile

Having modelled morphological nuclear phenotypes in *LMNA*-mutant muscle cells, we next sought to examine whether severely dysmorphic laminopathic myonuclei remain transcriptionally active and whether the transcriptional profile of the cells was altered compared to control cell lines. We first assessed the transcriptional status of *LMNA*-mutant myonuclei by labelling nascent RNA with the nucleoside analogue 5-ethynyl uridine (Figure 3A). Co-immunolabelling with eMyHC and Lamin A/C confirmed that all *LMNA*-mutant (including severely dysmorphic) myonuclei remain transcriptionally active (Figure 3B).

**Figure 3.**
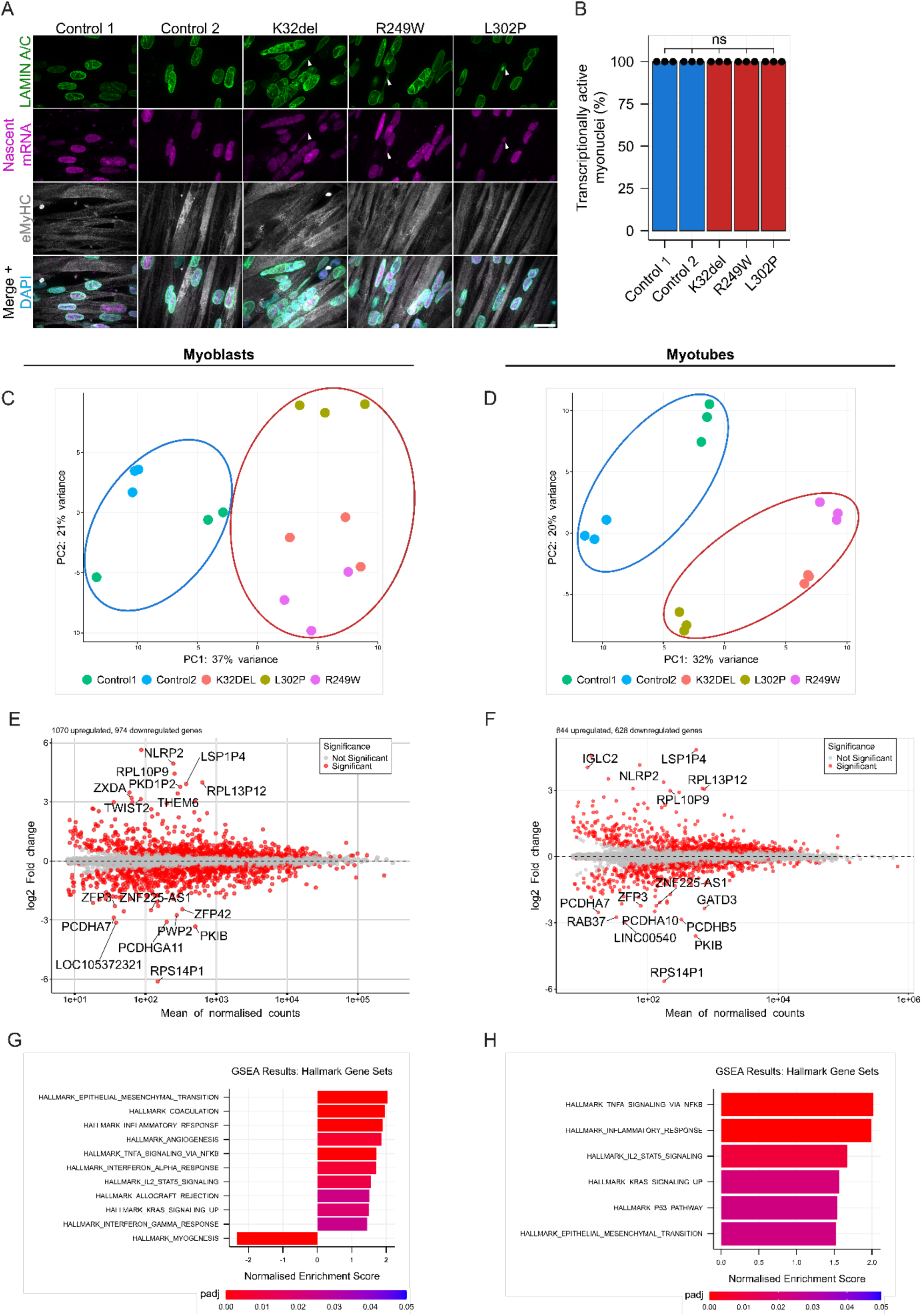
Transcriptomic profile of L-CMD iPSC-derived skeletal muscle cells. **(A)** Labelling of nascent RNA with alkyne-modified 5-ethynyl uridine and fluorescently tagged azide, co-immunostain of Lamin A/C and embryonic myosin heavy chain (eMyHC). Nuclei counterstained with DAPI. Arrows indicate dysmorphic myonuclei. Scale bar 20µm. **(B)** Quantification of transcriptionally active myonuclei. Fractional logistic regression with post-hoc pairwise comparisons using *Tukey’s* method. Three independent passages were analysed per cell line (N = 3). A total of 39-71 nuclei were analysed per cell line per repeat. ns p≥0.05. Dots represent the mean of one replicate. **(C, D)** PCA plots of control and *LMNA*-mutant (C) myoblasts and (D) myotubes. Blue circle indicates control cell lines and red circle indicates *LMNA*-mutant cell lines. **(E, F)** MA plots of *LMNA*- mutant versus control (E) myoblasts and (F) myotubes. Significantly differentially expressed genes are indicated in red while non-significant genes are indicated in grey. **(G, H)** Gene set enrichment analysis (GSEA) of *LMNA*-mutant versus control (G) myoblasts and (H) myotubes. Hallmark gene sets were used and significantly enriched gene sets are displayed as a bar chart ordered by normalised enrichment score (NES).

To examine transcriptional changes in *LMNA*-mutant iPSC derivatives, we performed bulk RNA sequencing of control and mutant myoblasts and myotubes. Following batch correction, we observed that control and *LMNA*-mutant samples clustered separately following principal component analysis (PCA) in both myoblast and myotube samples, though differences between control cell lines were present, presumably due to their different genetic background (Figure 3C, D).

Differential gene expression analysis revealed 1070 upregulated genes and 974 downregulated genes in *LMNA*-mutant myoblasts compared to controls (Figure 3E). For *LMNA*-mutant myotubes, we found 844 upregulated genes and 628 downregulated genes compared to controls (Figure 3F).

As two non-isogenic controls were used to examine transcriptional changes and these cell lines were transcriptionally distinct from each other in addition to being distinct from *LMNA*-mutants, we used gene set enrichment analysis (GSEA) to examine differential hallmark gene sets between *LMNA*-mutant and control cell lines. Notably, comparison of *LMNA*-mutant myoblasts to control lines revealed enrichment for hallmark gene sets related to inflammatory signalling including “Inflammatory response”, “TNFA signalling via NFκβ”, “Interferon alpha response” and “Interferon gamma response”. Such enriched gene sets indicate that cell autonomous inflammatory signalling may be altered in *LMNA*-mutant muscle cells potentially leading to activation of stress response pathways. Interestingly, “Myogenesis” was negatively enriched in *LMNA*-mutant myoblasts, despite our earlier observations that *LMNA*-mutant myogenic cells overtly differentiate as well as the controls (Figure 3G).

There were broadly similar transcriptional changes in *LMNA*-mutant myotubes when comparing to controls, including enrichment of the same inflammatory pathways “TNFα signalling *via* NFκβ” and “Inflammatory response”. Additionally, “P53 pathway” emerged as an enriched pathway in *LMNA*-mutant myotubes, indicating that these cells may be more transcriptionally primed for apoptotic cell death than controls (Figure 3H). Together, our results indicate that *LMNA*-mutant muscle cells have a cell-autonomous inflammatory transcriptional profile which may contribute to the severity of L-CMD pathophysiology.

### From disease to therapy modelling: L-CMD iPSC-derived myogenic cells to assess genetic therapies

Having established that the iPSC-based platform could model morphological and transcriptomic readouts in laminopathy skeletal myotubes, we sought to examine whether we could use our model system as a tool to development and test gene therapy strategies. Our previous work has shown that *LMNA* exon 3 and 5 are amenable to exon skipping, as removal of heptad repeats should not disrupt assembly of α-helical coils. This would be applicable to multiple patients sharing pathogenic variants within exon 5^8^. Removal of *LMNA* exon 5 did not cause an overt deleterious phenotype in primary murine embryonic fibroblasts, unlike exon 3-deletion^8^. Therefore, targeting a mutation in exon 5 would enable us to examine multiple modalities of gene therapy within our model system. To date, 243 individuals with mutations in *LMNA* exon 5 have been reported, with 52% of affected individuals having a striated muscle laminopathy phenotype (Figure 4A, B; www.umd.be/LMNA/, and G Bonne, R Ben Yaou, personal communication).

**Figure 4.**
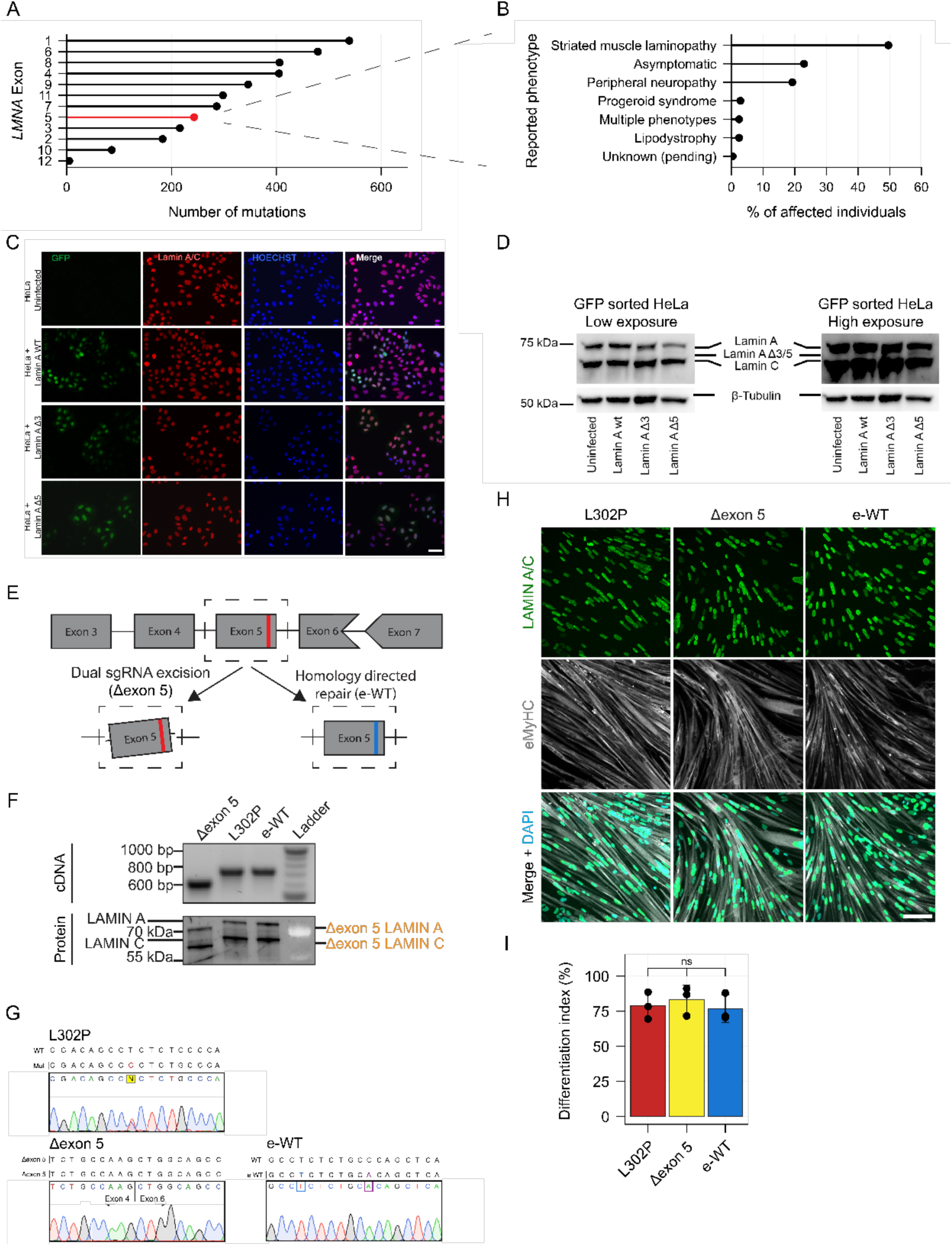
Two alternative CRISPR-based gene editing strategies targeting the *LMNA*^L302P^ variant in human iPSCs. **(A)** Graph showing the number of pathogenic variants reported in each *LMNA* exons in the UMD database (http://www.umd.be/LMNA/). **(B)** Graph showing the proportion of clinical phenotypes associated with pathogenic variants in *LMNA* exon 5 in UMD database. **(C)** Immunolabelling of HeLa cells transduced with Lamin A WT, Δexon 3 (as an additional control), or Δexon 5 constructs. GFP expressed through an IRES. Scale bar 100 µm. **(D)** Western blot of FACS sorted HeLa cells for Lamin A/C, showing the truncated Δexon 5 band at high exposure. **(E)** Schematic diagram of CRISPR editing strategies used, targeting the *LMNA*^L302P^ variant. **(F)** PCR of *LMNA* exon 5 region cDNA and western blot for Lamin A/C in *LMNA*^L302P^ and CRISPR edited cell lines. **(G)** Sanger sequencing chromatograms of *LMNA*^L302P^ and CRISPR edited cells. WT indicates wild type sequence context. L302P: Red letter is L-CMD-causing c.905T>C mutation; Δexon 5: junction of exons 4 and 6 is displayed; e-WT: Blue letter indicates C>T correction and purple letter indicates silent PAM-blocking mutation. **(H)** Immunostaining of LAMIN A/C and eMyHC. Nuclei counterstained with DAPI. Scale bar 100 µm. **(I)** Quantification of differentiation index. One-way ANOVA with Tukey’s post hoc comparisons to controls. Three independent passages were analysed per cell line (N = 3). 236-408 nuclei were analysed per cell line, per repeat. ns p ≥ 0.05. Dots represent mean value of one replicate. Whiskers show standard deviation.

We first used AlphaFold 2 to model predicted wild-type, L302P and exon 5-removed (Δexon 5) Lamin A and C protein dimers to predict the impact of exon removal on protein structure and dimerization. Homodimers of Δexon 5 Lamin proteins were predicted to assemble in a coiled-coil confirmation typical of Lamin dimers, whereas Δexon 5 and wild-type/L302P heterodimers were predicted to exhibit disrupted dimerization due to mismatched protein length (Figure S3A). We then investigated whether internally truncated lamins could be expressed, produced and assembled in human cells. HeLa cells were transduced with retroviral vectors overexpressing WT Lamin A, Lamin A Δexon 3 or Lamin A Δexon 5, all linked to a green fluorescent protein by an IRES site. Analysis of overexpressing cells showed correct production of the Δexon 3 and Δexon 5 internally truncated Lamin A proteins (Figure 4C, D), supporting further development of a genetic therapy strategy to skip or excise exon 5 using CRISPR. We also examined an alternative approach based upon precise correction of the c.905T>C, p.L302P L-CMD-causing missense mutation using CRISPR.

We targeted introns on both sides of exon 5 to excise it from the gDNA, thereby removing the mutation-containing exon entirely (Δexon 5) and in parallel, we corrected the *LMNA* L302P mutation in iPSCs using homology directed repair to generate edited wild-type (e-WT) iPSCs (Figure 4E; Figure S3B, C). We then differentiated these cells into myogenic precursors and examined formation of the expected RNA and protein products. e-WT cells produced cDNA and protein products of the same size as the L302P cells, while Δexon 5 cells showed shortened stable cDNA and protein species, corresponding to the shortened Lamin species produced (Figure 4F; Figure S3D, E). While generating clones of Δexon 5 lines, we also identified clones which had the expected gDNA band size, but abnormal cDNA and protein products which likely resulted from unintended DNA recombination events (Figure S3F, G). Therefore, these clones were ot used in further analysis. Sanger sequencing of cDNA of selected clones confirmed the expected genetic modifications (Figure 4G).

To examine whether CRISPR-based gene editing of the *LMNA* gene may impact differentiation capacity of the cells, we quantified differentiation index of the e-WT, L302P and Δexon 5 lines. Each of the cell lines produced multinucleated, myosin-positive myotubes as expected (Figure 4H), with similar differentiation index (Figure 4I; mean ±SD: L302P 78.82 ± 9.56 %, Δexon 5 82.23 ± 10.29 % and e-WT: 76.66 ± 9.85 %).

### L-CMD iPSC-derived skeletal myogenic cells enable testing CRISPR-based *LMNA* exon-removal and precise editing

Having generated L-CMD CRISPR-edited skeletal myogenic cells, we assessed whether established phenotypic readouts (e.g. Figure 2) would be ameliorated following gene editing. Examination of nuclear deformities revealed a significant amelioration of myonuclear shape phenotypes in corrected (edited) e-WT, but not Δexon 5 cells. The proportion of “Normal” myonuclei in the e-WT line was similar to that of Control 1 and Control 2 cell lines (Figure 2B), indicating a reversion of myonuclear shape phenotypes (mean±SD: L302P 64.05 ± 7.80 %; Δexon 5 74.06 ± 5.29 %; e-WT 87.45 ± 1.82 %; Figure 5B). Δexon 5 cells did not show a statistically significant restoration of nuclear morphological classifications when compared to L302P, and the proportion of “Deformed” myonuclei in L302P (30.24 ± 5.17 %) and Δexon 5 (22.72 ± 3.81 %) cell lines was higher than observed in e-WT myonuclei (11.53 ± 1.79 %). Similarly, “Blebbed” myonuclei were present at higher proportions in L302P (4.33 ± 2.66 %) and Δexon 5 (2.79 ± 1.25 %) than in e-WT (0.58 ± 0.26 %) cells. “Elongated” myonuclei were present in L302P (4.44 ± 4.62 %) and Δexon 5 (1.63 ± 1.01 %) cells but were completely absent from e-WT cells. Similarly, in keeping with our results in Figure 2B, L302P had a low proportion of “String” myonuclei (0.51 ± 0.5 %), as did Δexon 5 (0.43 ± 0.74 %), while these myonuclei were absent in e-WT cells (Figure 5B).

**Figure 5.**
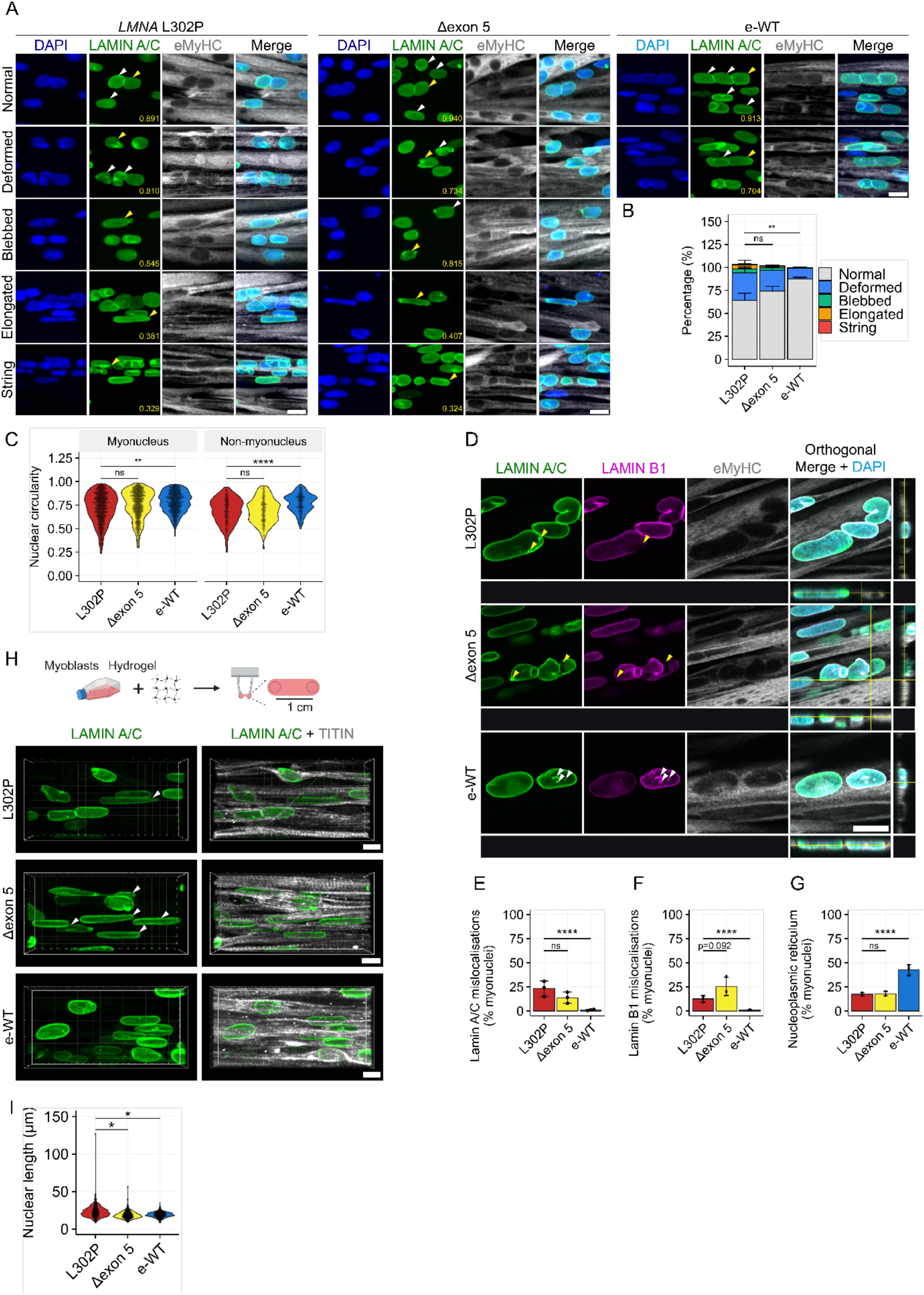
Phenotypic profile of CRISPR-edited *LMNA* L302P myonuclei. **(A)** Immunolabelling of Lamin A/C and embryonic myosin heavy chain (eMyHC) showing representative images of five nuclear morphology classifications are displayed: Normal, Deformed, Blebbed, Elongated, and String. Arrows indicate nuclei belonging to the indicated shape classification. Yellow arrows indicate nucleus associated with nuclear circularity measurement in lower right corner. Scale bar 20 µm. **(B)** Percentages of nuclear shape classifications in myotube nuclei. Nuclei with more than one type of abnormality (e.g. elongated and deformed) were scored twice meaning totals can exceed 100%. Nuclei were considered elongated if they had a major axis length >30 µm. Number of normal shaped nuclei in Δexon 5 and e-WT were compared to *LMNA* L302P for statistics. Fractional logistic regression with Tukey’s correction for pairwise comparisons. Three independent passages were analysed per cell line (N=3). 189 - 365 myotube nuclei were analysed per cell line, per repeat. ns p>0.05; ** p<0.01. Data presented as stacked bar chart. Whiskers show standard deviation. **(C)** Graphs showing quantification of nuclear circularity. Beta regression with Bonferroni correction for pairwise comparisons. Data presented as scatter plots showing measurements of all nuclei across repeats with violin plots showing distribution of measurements. Three independent passages were analysed per cell line (N=3). 181 - 301 myotube nuclei and 32 - 108 non-myotube nuclei were analysed per cell line, per repeat. ns p≥0.05; ** p<0.01; **** p<0.0001. **(D)** Representative immunolabelling of Lamin A/C, Lamin B1 and eMyHC. Counterstained with DAPI. Z-stack images of control and LMNA-mutant nuclei are displayed as a single Z-slice, with orthogonal view. White arrowheads indicate nucleoplasmic reticulum, yellow arrows indicate Lamin mislocalisations. Scale bar 20 µm. **(E, F)** Lamin A/C (E) and Lamin B1 (F) mislocalisation. Fractional logistic regression with Tukey’s post hoc test for pairwise comparisons. Three independent passages were analysed per cell line (N = 3). A total of 237 - 340 nuclei were analysed per cell line, per repeat. ns p≥0.05; **** p < 0.0001. Dots represent the mean value of one replicate. Whiskers indicate the standard deviation. **(G)** Quantification of nucleoplasmic reticulum positive nuclei in LMNA-mutant, Δexon5 and e-WT myotube nuclei. Fractional logistic regression with Tukey’s post hoc test for pairwise comparisons. Three independent passages were analysed per cell line (N = 3). 237 - 340 nuclei were analysed per cell line, per repeat. ns p≥0.05; **** p<0.0001. Dots show mean value of one replicate. Whiskers show standard deviation. **(H)** Schematic diagram showing generation of 3D engineered muscles. Immunostaining for Lamin A/C and Titin in 3D engineered muscles. Arrows indicate elongated myonuclei. Scale bar 5 µm. **(I)** Graph showing quantification of myonuclear length. One-way ANOVA with Tukey’s correction for pairwise comparisons on log-normalised data to achieve normal distribution. Data presented as scatter plots showing measurements of all nuclei across repeats with violin plots showing distribution of measurements. Three independent passages were analysed per cell line (N=3). 88 - 357 were analysed per cell line, per repeat. * p<0.05.

For an objective readout, the NCR was quantified comparing Δexon 5 and e-WT nuclei to *LMNA* L302P nuclei (Figure 5C). L302P myonuclei and non-myonuclei showed reduced circularity (myonuclei: 0.727 ± 0.063; non-myonuclei: 0.676 ± 0.057). Δexon 5 myonuclei did not achieve significant improvement of circularity scores (myonuclei: 0.760 ± 0.098; non-myonuclei: 0.692 ± 0.08). In contrast, precise correction of the L302P mutation in the e-WT line significantly ameliorated NCR in both myonuclei and non-myonuclei (myonuclei: 0.781 ± 0.044; non-myonuclei: 0.781 ± 0.054). Analysis of myonuclear perimeter showed no significant difference between L302P and either Δexon 5 or e-WT, whereas perimeter of e-WT non-myonuclei (56.58 ± 0.78 µm) were significantly shorter versus L302P (63.21 ± 3.49 µm), while no significant amelioration was observed in Δexon 5 (60.31 ± 4.76 µm). Furthermore, nuclear area did not differ significantly in any of the groups; Figure S4.

We then analysed lamin mislocalisation in CRISPR-edited cells (Figure 5D). Similar to nuclear shape analyses, there was no statistically significant amelioration of Lamin A/C mislocalisation compared to L302P (23.2 ± 8.04 %) in Δexon 5 myonuclei (13.8 ± 5.75 %; Figure 5E). In contrast, e-WT myonuclei showed significant amelioration of Lamin A/C mislocalisation (1.07 ± 0.576 %). Mean Lamin B1 mislocalisation was unchanged in Δexon 5 myonuclei (25.4 ± 9.40 %) compared to L302P myonuclei (12.5 ± 3.43 %; Figure 5F). In contrast, e-WT myonuclei Lamin B1 mislocalisation was significantly improved (1.05 ± 0.50 %). Finally, we analysed the formation of nucleoplasmic reticula in the myonuclei of L302P and CRISPR-corrected cells (Figure 5G). Similar to other analyses, nucleoplasmic reticulum formation was not ameliorated in Δexon 5 myonuclei (17.86 ± 2.37 %) compared to L302P (17.35 ± 1.56 %). Myonuclei positive for nucleoplasmic reticulum were significantly increased in e-WT (42.69 ± 5.68 %) myonuclei, indicating that these structures are stabilised in cells expressing wild type Lamin A/C.

Having established that e-WT cells ameliorate characteristic myonuclear morphological and protein localisation, we next utilised the 3D engineered muscle platform to examine amelioration of myonuclear phenotypes in a physiologically relevant culture system (Figure S4C). Analysis of myonuclear shape in engineered muscles showed that both Δexon 5 (18.72 ± 0.16 %) and e-WT (18.79 ± 0.99 %) nuclei were significantly less elongated than L302P (22.86 ± 2.56 %), demonstrating amelioration of nuclear elongation phenotypes with CRISPR-based interventions in the 3D culture system (Figure 5H).

### Comparative assessment of two CRISPR-Cas9 strategies shows that precise *LMNA* editing, but not exon-removal, ameliorates transcriptomic signature

Having established that our iPSC-based skeletal myogenic platform had sufficient resolution to quantify and compare amelioration of morphological phenotypes in *LMNA* L302P cell lines treated with different gene editing strategies, we asked whether the same platform could detect changes in the transcriptomic profile of edited cells. We also hypothesised that differentiating myogenic cells in 3D engineered muscles may increase transcriptional differences between groups due to enhanced maturation of myofibres associated with the enhanced physiological relevance of the system (e.g., cell-cell and cell-matrix 3D interactions, increased mechanical tension).

We therefore induced skeletal myogenesis in *LMNA* L302P, Δexon 5 and e-WT cell lines and performed bulk RNA-sequencing in 2D and 3D cultures at sequential stages of differentiation: myoblasts, myotubes in 2D (differentiated for 4 days), and myotubes in 3D (differentiated for either 4 or 7 days). PCA1 (80.1% variance) distinguished myoblast, 2D myotube and 3D engineered muscle samples, demonstrating a transcriptional shift between cell states and culture systems (Figure 6A). We then examined whether there was a difference in expression of myogenic genes between groups (Figure S5A). All myotubes showed upregulation of sarcomeric genes compared to myoblasts. However, compared to 2D myotubes, 3D engineered muscles at a time-matched 4 days of differentiation or 7 days of differentiation showed downregulation of *MYH3*, an embryonic myosin associated with early development and upregulation of *MYH8*, a myosin associated with secondary (foetal/perinatal) myogenesis. Additionally, transcripts for the giant proteins Titin (*TTN*) and Nebulin (*NEB*) were upregulated in 3D engineered muscles compared to 2D myotubes, validating that engineered muscle platform enhances maturation of myofibres (Figure S5A).

**Figure 6.**
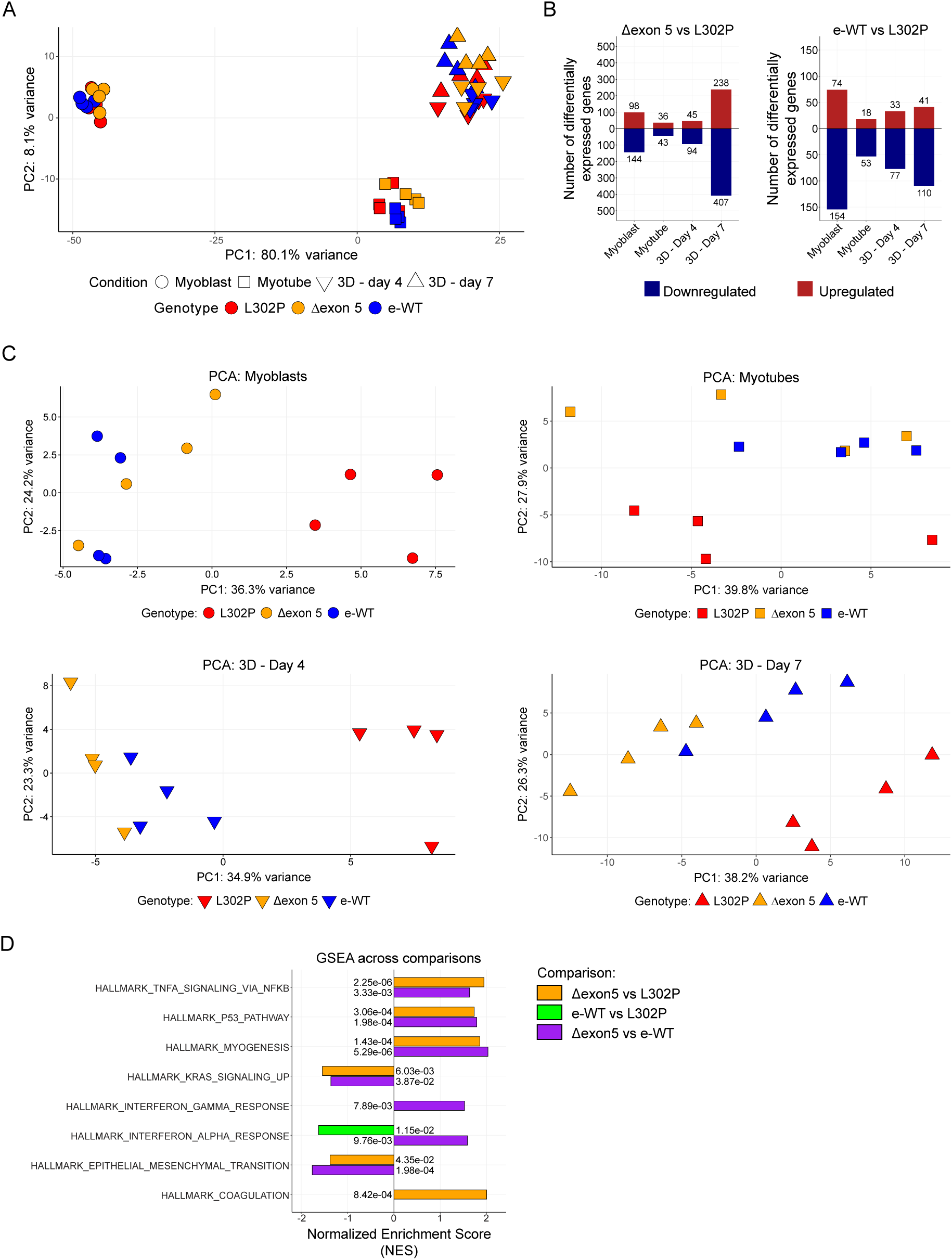
Transcriptomic profile of *LMNA* L302P and CRISPR-edited myogenic cells across multiple cell states and culture conditions. **(A)** PCA of myoblasts, and differentiated 2D myotubes and 3D engineered muscles from *LMNA* L302P, Δexon 5 and e-WT lines. **(B)** Diverging bar chart showing number of differentially expressed genes for Δexon 5 and e-WT vs L302P in each culture condition. **(C)** individual PCA plots for myoblasts, 2D myotubes, 3D engineered muscles differentiated for 4 days and 3D engineered muscles differentiated for 7 days. **(D)** GSEA of Δexon 5 vs L302P, e-WT vs L302P and Δexon 5 vs e-WT comparisons in 3D engineered muscles at day 7 of differentiation. Indicated hallmark gene sets were used for analysis. All significantly enriched gene sets (padj < 0.05) in each comparison are shown. Numbers show padj value for adjacent gene set and comparison.

We next examined transcriptional differences between *LMNA* L302P cells and CRISPR edited Δexon 5 and e-WT cells within each condition. Analysis of differentially expressed genes revealed distinct patterns, with varying numbers of differentially expressed genes between Δexon 5 and e-WT cells compared to L302P (Figure 6B). Myoblast comparisons revealed 98 upregulated and 144 downregulated genes in Δexon 5 versus L302P, and 74 upregulated and 154 downregulated genes in e-WT versus L302P. Consistent with our previous analysis (Figure 3), the number of differentially expressed genes was reduced in myotubes compared to myoblasts. Comparison of myotubes showed 36 upregulated and 43 downregulated genes in Δexon 5 versus L302P, and 18 upregulated and 53 downregulated genes in e-WT versus L302P. Engineered muscles showed an increase of differentially expressed genes in a time dependent-manner compared to myotubes. At 4 days of differentiation, 45 genes were upregulated and 94 genes downregulated in Δexon 5 versus L302P, while at 7 days of differentiation, this number increased to 238 upregulated and 407 downregulated genes versus L302P. Similarly, an increasing pattern of differentially expressed genes was observed in the e-WT versus L302P comparison with 33 upregulated and 77 downregulated genes at 4 days of differentiation, and 41 upregulated genes and 110 downregulated genes at 7 days of differentiation.

Given our observation that transcription of late-stage muscle genes such as *TTN* and *NEB* are increased in 3D engineered muscles compared to 2D myotubes, we hypothesised that the increasing number of differentially expressed genes across conditions may be driven by two factors: 1. increased myogenic maturation across cell culture systems and time, and 2. prolonged exposure to mutant Lamin A/C proteins. We therefore investigated transcriptional similarity of genotypes across conditions (Figure 6C). In myoblasts and 2D myotubes, L302P clustered away from Δexon 5 and e-WT samples, indicating early transcriptional changes in both conditions. In 3D engineered muscles at both 4 and 7 days of differentiation, L302P again clustered away from Δexon 5 and e-WT samples. However, a separation between Δexon 5 and e-WT samples began to emerge at 4 days of differentiation and became evident at 7 days of differentiation, suggesting that while Δexon 5 cells are initially transcriptionally similar to e-WT cells, the presence of myonuclear deformities may lead to the emergence of a “laminopathic” transcriptional profile.

To further assess changes in transcriptional profiles of CRISPR-treated cells, we applied GSEA comparing Δexon 5 and e-WT cells in each condition to L302P. Comparisons of Δexon 5 samples to L302P, across myoblast, myotube and engineered muscles surprisingly showed enrichment for pro-inflammatory and apoptotic pathways such as “TNFα signalling *via* NFKβ”, “p53 signalling” and “Interferon-α response” (Figure S5B). Consistent with analysis of differentially expressed genes (Figure 6B), we observed the greatest number of enriched gene sets in engineered muscles differentiated for 7 days (Figure 6D). Interestingly, this comparison showed upregulation of the “Coagulation” gene set in Δexon 5 cells, similarly to that reported in conditions linked to pro-inflammatory signals such as myocardial infarction, osteoarthritis and colorectal cancer^39–41^. In combination with a lack of amelioration of morphological myonuclear abnormalities (Figure 5), these results indicate that removal of *LMNA* exon 5 in human myogenic cells might potentiate rather than ameliorate laminopathy disease phenotypes.

Conversely, examination of enriched gene sets in e-WT cells *versus* L302P displayed relatively fewer enriched pathways (Figure S5C). There were no enriched terms in 3D engineered muscles at day 4 of differentiation, indicating that this may be too early a time-point to discern transcriptional differences in isogenic experiments such as these. Encouragingly, we observed a downregulation of the pro-inflammatory gene set “Interferon alpha signalling” in e-WT cells at 7 days of differentiation in e-WT 3D engineered muscles, indicating that an amelioration of transcriptional pro-inflammatory signals started to emerge using this model system (Figure 6D).

Finally, we compared the effect of *LMNA* exon 5 removal against precise gene editing at 7 days of differentiation in engineered muscles, to examine if Δexon 5 cells were enriched for pro-inflammatory gene sets against another CRISPR-treated line. We observed enrichment in Δexon 5 cells for gene sets including “p53 pathway”, “TNFα signalling *via* NFκβ”, “Interferon-α response” and “Interferon-γ response” (Figure 6D). These results confirm that the pro-inflammatory profile is not caused by CRISPR treatment of the *LMNA* gene but rather is associated with *LMNA* exon 5 deletion.

Overall, these transcriptomic data support the conclusion that in human skeletal myogenic cells, *LMNA* exon skipping may not a viable therapeutic strategy due a failure to ameliorate phenotypic and transcriptional (pro-inflammatory) phenotypes. In contrast, combining iPSC-based modelling with advanced 3D engineered muscle model systems demonstrates that precise genetic correction of *LMNA* variants associated with L-CMD using CRISPS-Cas9 gene editing leads amelioration of multiple transcriptional and phenotypic readouts. In addition, this validates the use of these laminopathy disease associated phenotypes as reliable readouts for use in therapy screening platforms.

## DISCUSSION

Models of skeletal muscle laminopathies have relied primarily on platforms which do not recapitulate human genetics (e.g., mouse, *Drosophila*), are not in the context of the primarily affected tissue (e.g., fibroblasts, HEK or HeLa cell lines) or are difficult to access (e.g., human biopsies). Immortalised myoblasts address some of those limitations, but do not allow studies into early pathogenic events (i.e. during developmental myogenic specification). Use of iPSC-based models can address most of these challenges and in this study, we focused on harnessing their potential to gain insights into the pathogenesis of laminopathies, as well as to model response to genetic therapies.

Previous studies showed impaired myogenic differentiation in *Lmna*-mutant myoblasts^23,31,32^. Overexpression of *LMNA* mutations W520S or R453W in C2C12 cells,^33,42^ K32del mouse primary myoblasts and K32del, R249W and L380S human immortalised myoblasts all displayed impaired myogenic differentiation^5,28,43^. Transcriptional and epigenetic mechanisms leading to expression of non-muscle genes have been suggested as causative mechanisms^20^. However, most studies focus on murine myogenesis and those based upon human cells likely model adult/regenerative myogenesis. iPSCs derived from patients with several congenital laminopathy phenotypes allowed us to examine whether perturbations of myogenesis occurred at earlier developmental stages. Exogenous myogenic factors were not used in hiPSCs to avoid confounding factors that may compensate for any myogenic defect as in our previous studies^9,12^. Using directed differentiation of iPSCs as a synthetic model of human developmental myogenesis, we did not find overt defects in myogenesis. Thus, defects in myogenic differentiation and loss of muscle-specific gene expression could be more prevalent postnatally or require prolonged exposure of the cell to an abnormal nuclear lamina and its consequences, including abnormal chromatin configuration, DNA damage accumulation or nuclear ruptures.

Nuclear shape abnormalities are an established phenotype in striated muscle laminopathies reported *in vivo* and *in vitro*^4–9,9,10,44–46^. The K32del and R249W variants used here display nuclear shape abnormalities^4,5,47–49^, and are associated with significant clinical severity^50^. Abnormal nuclear morphometric parameters were also present in a novel line harbouring the *LMNA* L302P variant. Thus, our model recapitulated nuclear morphological abnormalities in all *LMNA*-mutant lines assessed, including nuclear elongation in 3D culture, demonstrating it as a robust platform for modelling disease-associated pathological features.

Lamin mislocalisation contributes to weakening of the nuclear envelope and is associated with nuclear rupture^14,44,51^. Additionally, lamin mislocalisation and aggregation are predictive of affected tissue type with most myopathic *LMNA* mutations showing aggregation^36^. Those findings were reproduced in our platform and, interestingly, iPSC-based disease modelling using developmentally relevant approaches for myogenic differentiation resulted in better capture of lamin protein mislocalisation than transgene-based approaches^9^. For example, Lamin A/C and B1 mislocalisation with *LMNA* K32del and R249W was not evident using a transgene-based iPSC differentiation protocol^9,52^.

There was reduced formation of nucleoplasmic reticulum in *LMNA*-mutant myonuclei. These nuclear invaginations have reported roles in maintenance of nuclear architecture, nucleocytoplasmic transport, calcium signalling and DNA damage repair^37,53^. Accumulation of progerin at the nuclear envelope in Hutchinson-Gilford progeria syndrome (HGPS) increases the prevalence of nucleoplasmic reticulum in HGPS nuclei^54^. Furthermore, the EDMD-causing R453W and the FPLD2-causing R482W mutations also lead to an increase in nucleoplasmic reticulum in fibroblasts. In contrast, our observation that nucleoplasmic reticula are reduced in *LMNA*-mutant myogenic cell nuclei suggests that these intranuclear structures might be destabilised by L-CMD pathogenic variants, though further examination of these changes in patient cells of other lineages is needed to determine if this is a cell type-specific effect.

An outstanding question is how changes in nuclear architecture affect transcriptional output. We found that dysmorphic nuclei remain transcriptionally active even in cases of severe deformity. This highlights the need for closer examination of chromatin organisation and transcriptional output in structurally compromised myonuclei. Our assessment of bulk transcriptional output to ascertain global changes in transcription in *LMNA*-mutant myogenic cells identified multiple hallmark gene sets related to inflammatory signalling were enriched in *LMNA*-mutant iPSC skeletal myogenic derivatives. This is consistent with observations in mouse *Lmna*-null and *Lmna*^D300N^ cardiac tissues^55,56^, and in muscle biopsies of patients with skeletal muscle laminopathies, where an atypically florid inflammatory infiltrate and fibrosis has been reported^2^. Moreover, this is also in keeping with high levels of proinflammatory and profibrotic cytokines reported in the serum of patients with muscle laminopathies, pointing to a cell autonomous source (i.e. muscle fibres) for some of those signals^57,58^. Interestingly, case-reports of beneficial corticosteroid therapy in some children with L-CMD would be in keeping with this observation, supporting further investigations into this strategy as a possible therapeutic option targeting the inflammatory component of the disease^59^.

A robust *in vitro* disease model should have sufficient resolution to predict outcomes of possible therapeutic interventions. We previously reported a potential exon-skipping strategy for pathogenic variants in *LMNA* exon 5, conducted primarily in primary mouse embryonic fibroblasts (pMEFs)^8^; however, phenotypes differ between mouse and human^26^. Therefore, we used our human model to examine exon-skipping and precise correction/editing as gene therapy strategies for L-CMD. Excision of *LMNA* exon 5 lead to Lamin A/C production, nuclear envelope assembly and myogenic differentiation, but with no overall amelioration of myonuclear defects. Precise CRISPR-correction of the *LMNA*^L302P^ variant reverted myonuclear morphological phenotypes to a state comparable with healthy controls, supporting further pre-clinical work to assess delivery modalities and gene editors capable of targeting and correcting pathogenic *LMNA* variants also in post-mitotic nuclei.

Beyond morphological parameters, transcriptional output regulates cell and tissue behaviour. Therefore, a therapeutic approach which may normalise some or all morphological phenotypes, but not transcriptomic output may not ultimately provide therapeutic benefit. Examination of transcriptional output demonstrated that Δexon 5 myogenic cells were enriched for multiple pro-inflammatory gene sets compared to the parental *LMNA* L302P line, establishing that restoration of some nuclear morphological parameters alone is not sufficient to restore cellular homeostasis. In contrast, our 3D engineered muscle model demonstrated that precise correction of the *LMNA*^L302P^ variant via CRISPR-Cas editing resulted in downregulation of the pro-inflammatory “Interferon-α response” gene set. consistent with previous work showing activation of pro-inflammatory signalling cascades upon mutation or knockout of Lamin A/C, associated with DNA damage and/or DNA leakage into cytoplasm sensed by the cGAS/STING pathway^16,25,44,55,57,60,61^. Our study provides evidence that this phenomenon occurs remarkably early in L-CMD pathophysiology, as it is detectable in culture systems representative of developing muscle (likely foetal-perinatal, based on our transcriptional profiling) and exposed to mutant lamins for a relatively short timeframe (as they were not produced in the pluripotent iPSC colonies).

iPSC reprogramming of *LMNA*-mutant cells can reverse HGPS phenotypes, which are then reacquired upon differentiation^62^. This is likely due to Lamin A/C expression being low to absent in pluripotent cells and upregulated upon differentiation. Here, we applied gene therapy approaches directly to iPSCs which we then differentiated into myogenic cells. Future work should examine the impact of similar gene therapies on cells which have a nuclear lamina containing Lamin A/C, such as post-mitotic muscle tissue. It is of note that Lamin A/C proteins exhibit a long half-life^63^ and so when gene therapy is applied to non-dividing cells, it may take some time for the newly repaired Lamin A/C proteins to have an ameliorating effect on disease phenotypes^63^.

Overall, here we provide evidence that hiPSCs and their skeletal myogenic derivatives can contribute to several key aspects of translational research in laminopathies, including: 1) diagnostics, such as detection of nuclear abnormalities in novel gene variants; 2) pathophysiology, including assessment of myogenic defects and proinflammatory signature; and 3) therapy development, via testing CRISPR-based gene editors. Together, this advances complex disease modelling in rare diseases.

## MATERIALS AND METHODS

### Cell lines and cultures

Two healthy control human iPSC lines were used: NCRM1 (https://hpscreg.eu/cell-line/CRMi003-A) and NCRM5 (https://hpscreg.eu/cell-line/CRMi001-A), referred to as Control 1 and 2 respectively. *LMNA* K32del (c.94_96delAAG; https://hpscreg.eu/cell-line/CMDi003-A) and R249W (c.745C>T; https://hpscreg.eu/cell-line/CMDi004-A) cell lines were kindly provided by Cellular Dynamics International Inc. (https://www.fujifilmcdi.com/) and Cure Congenital Muscular Dystrophy (CureCMD: https://www.curecmd.org/). iPSCs were generated by CDI by reprogramming samples provided by CureCMD (holder of clinical information) using episomal vectors. *LMNA* L302P iPSCs were generated by lentiviral expression of pluripotency genes (c.905T>C; https://hpscreg.eu/cell-line/UCLi006-A; see below for details).

iPSCs were cultured in mTeSR Plus medium (STEMCELL Technologies, 100-0276), supplemented with 100 IU penicillin/streptomycin. Cells were grown on tissue culture plates coated in Matrigel (Corning, 354230/354231), diluted 1:100 in DMEM high glucose at 5% O_2_ and 5% CO_2_. Skeletal myogenic differentiation was performed using Myocea skeletal muscle differentiation kit (Myocea Inc., SKM01, SKM02) ^34^. Briefly, iPSCs were dissociated in TrypLE Express (GIBCO, 12604-013) and 5,000 cells/cm^2^ were seeded into matrigel coated plates. Cells were maintained in induction medium (SKM01) for 10 days, with medium changes every other day. At day 10, cells were passaged and plated on Matrigel coated plates at 5000 cells/cm^2^ in Myoblast medium (SKM02). Cells were maintained for 8 days, with medium changes every other day. At day 8, cells were passaged and from this point, were maintained and passaged in tissue culture flasks without Matrigel in Myoblast medium. To differentiate myoblasts into multinucleated myotubes, myoblasts were passaged and plated on Matrigel coated plates at ∼60,000 cells/cm^2^ at a confluence of 95-100% after attachment. The next day, cells were switched to differentiation medium (DMEM high glucose (SIGMA, D5671-500mL), 2% horse serum (Euroclone), 1% Glutamax (Thermo Fisher Scientific, A5256701), 1% penicillin/streptomycin (GIBCO, 15070-063). In monlayer cultures, cells were differentiated for 3-4 days. In 3D engineered muscles, cells were differentiated for 5-7 days with 3 parts differentiation medium, 1 part Myoblast medium.

HeLa cells were cultured in DMEM High glucose (Sigma, D5671-500mL), supplemented with 10% FBS (Life Technologies, A5670701), 2 mM Glutamax (Sigma, 35050061), 100 IU + 0.1 mg/mL penicillin/streptomycin (Sigma, P4333-100ML). Cells were grown at 37°C under ambient O_2_ and 5% CO_2._ Human cell work was conducted under the approval of the NHS Health Research Authority Research Ethics Committee reference no. 13/LO/1826; IRAS project ID no. 141100.

### Generation and characterisation of *LMNA*^L302P^ hiPSCs

#### Reprogramming to pluripotency

*LMNA*^L302P^ dermal fibroblasts were kindly provided by Jean-Marie Cuisset (Centre de Référence des maladies rares neuromusculaires Nord/Est/Ile de France, Lille, France) under informed consent and reprogrammed using lentiviral vectors encoding MYC, SOX2, OCT3/4, and KLF4 under the control of CMV promoters (Kerafast) and Human iPSC Boost Supplement II kit (Merck Millipore, SCM094). Reprogramming was conducted on mitotically inactivated mouse embryonic feeder (MEF) based culture conditions with Knockout serum replacement medium (Knockout DMEM (Gibco, 10829018), 25% (v/v) KSR (Gibco, 10828028), 1% non-essential amino acids (Sigma, M7145-100ML), 2 mM Glutamax (Sigma, 35050061), 0.1 mM β-mercaptoethanol (Gibco, 21985023), 100 IU + 0.1 mg/mL penicillin/streptomycin (Sigma, P4333-100ML), 10 ng/mL bFGF (Gibco, 13256-029)) according to manufacturer’s recommendations. 80,000 cells were plated in 6 well culture plates. Cells were transduced with the four viruses at MOI 30 in 1 mL of 10% DMEM (Dulbecco’s modified eagle medium) with 8 μg/mL Polybrene (hexadimethrine bromide, Sigma). For the viral transductions, heat inactivated DMEM 10% was used (FBS heat inactivated at 56 °C for 30 minutes, to inactivate complement factors). Cells were incubated with the viral mix for 8 hours, after which the medium was aspirated, cells washed three times with PBS, and fresh DMEM 10% added. On day 5 transduced cells were replated onto Matrigel coated plates on top of a layer of inactivated MEFs, with knockout serum replacement medium supplemented with Human iPSC Boost Supplement II kit. Two six well plates of MEFs were used to plate infected fibroblasts. Medium was changed every other day until day 25. Fresh MEFs were added to the wells on day 11. Small molecule supplementation was added to the media every other day until day 18. On day 25 the first iPSC colonies were isolated and picked manually to isolate from fibroblasts that had not been reprogrammed. Isolated colonies were plated onto freshly prepared plates of MEFs. Each picked colony was cultured separately, to ensure clonal expansion. Three distinct clonal lines were expanded in knockout serum replacement medium on MEFs for ∼4 passages, after which they were moved to feeder-free conditions with TESR-E8 medium and expanded on Matrigel following standard manufacturer’s recommendations (Stem Cell Technologies).

#### Karyotype analysis

Karyotype analysis was performed by The Doctors Laboratory (TDL, London, UK). For *LMNA* L302P iPSCs, 7 cells were analysed. 5 showed a normal karyotype, with 2 having chromosome losses associated with preparation artefacts.

#### Alkaline phosphatase staining

Cells were washed with PBS, fixed with 4% PFA for 5 minutes and washed again with PBS. Cells were incubated for 20 minutes in alkaline phosphatase staining solution (0.1 M Trizma hydrochloride, 0.1 M NaCl, 0.5 M MgCl_2_, 0.5 % v/v µL Tween 20, 2 % v/v NBT/BCIP solution (18.75 mg/mL NBT, 9.4 mg/mL BCIP). Following incubation, cells were imaged with a dissecting microscope.

#### Embryoid body formation

iPSCs were detached with Gentle Cell Dissociation Reagent for 6 minutes at room temperature, after which it was aspirated, and TESR-E8 containing 4 mg/mL of Polyvinal alcohol and 3 μg/mL of ROCK inhibitor Y27362 (Calbiochem) was added. Cells were collected by scraping and transferred to a non-tissue culture treated plate, to allow EBs to form in suspension. The following day the medium was changed to TESR-E6 (Stem Cell Technology), for 5 days, with media changes every other day. After five days the medium was switched to spontaneous differentiation medium (DMEM 20 %), although cells were not transferred to adherent plates (tissue culture treated), the EBs adhered to the non-tissue culture treated plate after the medium was switched to DMEM 20%. After 7 days, the medium was aspirated, replaced with PBS and cells collected by scraping, cells were spun at 232 g, transferred to a 1 mL Eppendorf vial and spun again at 10,000 g. RNA was then extracted from the pellet, as described below..

### 3D engineered skeletal muscle constructs

3D engineered muscles were constructed as previously described^12,64^. Briefly, myoblasts were pre-treated with 10 µm ROCK inhibitor for 2 hours prior to passaging. Cells were counted (1 million/construct) and centrifuged before being placed on ice. 100 µL of heat-inactivated Myoblast medium was added, after which, 10% Matrigel and 3.5 mg/mL human fibrinogen (Baxter) were added. Inactivated medium was added to the calculated final volume and for each muscle, 113uL was individually added to aliquoted thrombin (Baxter). The final mixture for each muscle was decanted into 2% UltraPure agarose in DPBS (Invitrogen, 16500-500) moulds containing silicon posts (EHT Technologies, GmbH Hamburg). Constructs were placed at 37°C, 5% CO2 for 2 hours before DMEM high glucose was added onto each muscle. After a further 10-minute incubation at 37°C, posts were removed from the moulds and muscles were visually inspected for successful polymerisation. Muscles were placed into a fresh 24 well plate containing pre-warmed Myoblast medium supplemented with 33 µg/µL aprotinin (Sigma Aldrich, A3428-10MG). Muscle constructs were cultured for 2 days in Myoblast medium at 37°C, 5% CO_2_ after which the medium was switched to differentiation medium.

### Generation of CRISPR-corrected *LMNA*^L302P^ iPSCs (e-WT)

CRISPR correction of *LMNA* L302P iPSCs was performed by Synthego. *LMNA* L302P iPSCs were sent as frozen stocks and homology directed repair was performed. sgRNA and donor template used are indicated in Supplementary table 1.

#### CRISPR sgRNA and pegRNA design

sgRNAs were designed using Benchling CRISPR design tool using cutting efficiency calculations based on Doencsh *et al*.^65^ sgRNAs were ordered from Synthego with 2’-O-Methyl analogues and 3’ phosphorothioate bond modifications. sgRNAs used are indicated in Supplementary table 2.

#### Nucleofection

Nucleofection of iPSCs was performed using an Amaxa 4D nucleofector X unit (Lonza, AAF-1003X) and the P3 Primary Cell Kit (Lonza, V4XP-3032). Synthego sgRNAs were hydrated to 100 µM in TE buffer. 2 hours prior to nucleofection, iPSCs were treated with a final concentration of 10 µM ROCK inhibitor (Cell Guidance Systems, SM02-5). 1 hour prior to nucleofection, 6 well plates were coated with Matrigel. For excision of *LMNA* exon 5 sequence, 1.5 µL of each sgRNA was premixed with 2 µL 20 µM SpyCas9 (NEB, M0646T) to a final concentration of 300 pmol gRNA (150 pmol each) and 40 pmol SpyCas9 respectively. The mixture was incubated at room temperature for 10 minutes and was transferred to ice. Cells were detached with TrypLE Express for 5 minutes and neutralised with premade mTesr1 + CloneR2 (STEMCELL Technologies, 100-0691). 250,000 cells were counted per reaction and transferred to separate 1.5 mL Eppendorf tubes. Cells were resuspended in 20 µL P3 primary solution and mixed with pre-made RNP complex. The mixture was transferred to nucleofector cuvettes and nucleofected using program CA137. Pre-warmed mTesr1 + CloneR2 was added to the cuvette and incubated for 5 minutes before cells were transferred dropwise to 6 well cell culture dishes with pre-warmed mTesr1 + CloneR2. Cells were maintained for 3 days before being dissociated using TrypLE Express. Half of the cells were taken for cryopreservation, and the other half was used for DNA extraction and PCR. Once genome editing had been confirmed by PCR, cells were thawed and expanded using standard iPSC culture techniques.

#### Single cell cloning

Single cell sorting of iPSCs was carried out in the Flow Cytometry facility at the Francis Crick Institute on a FACS ARIA II. Prior to dissociation, cells were treated for 2 hours with 10 µM ROCK inhibitor. Cells were dissociated with TrypLE Express for 3-5 minutes at 37°C, centrifuged and resuspended in 1 mL of mTesr1 + CloneR2 (STEMCELL Technologies, 100-0691). Cells were passed through a FACS tube filter and were transported for sorting. Cells were cloned into 96 well plates pre-coated with Matrigel and filled with mTesr1 + CloneR2 using a 100µm nozzle. Cells were subsequently fed every two days with mTesr1 + CloneR2 thereafter. Cells were monitored regularly and approximately 1 to 2 weeks later, surviving colonies were expanded to a 24 well and then 12 well plate format in this medium using standard iPSC culture techniques. Once ready to passage in a 12 well plate, half the cells were cryopreserved, and half were taken for DNA isolation and PCR-based screening.

### DNA / RNA extraction and cDNA preparation

DNA was extracted from cells using DNAeasy blood and tissue kit (QIAGEN, 69506) as described in the protocol. DNA was eluted in elution buffer (EB). DNA concentration and purity was verified by nanodrop. RNA extraction was performed using RNAeasy Micro Kit (QIAGEN, 74004) as described in the protocol, with the exception that all centrifugation times were doubled. RNA was eluted in RNase-free water and quantified by Nanodrop. cDNA was prepared from extracted DNA using High-Capacity RNA-to-cDNA Kit (Applied Biosystems, 4387406) as described in the protocol.

### Sanger sequencing

PCR for Sanger sequencing was carried out using Promega GoTaq G2 Hot Start Taq Polymerase (Promega, M7422) according to the manufacturer’s protocol. Samples were purified using GeneJet PCR Purification Kit (Thermo Fisher Scientific, K0701). Sanger sequencing was performed by GENEWIZ LLC. Primer sequences are provided in Supplementary Table 3.

### RNA-sequencing

RNA was extracted for RNA sequencing using RNAeasy Micro Kit (QIAGEN, 74004). Extracted RNA was quantified using a Bioanalyzer. RNAseq libraries were prepared using NEBNext Ultra II Directional PolyA mRNA kit within the Genomics facility at The Francis Crick Institute. Sequencing was performed in paired-end 100bp on Novaseq S2 platform in the Genomics facility in The Francis Crick Institute to a depth of 25 million reads per sample. RNA sequencing data was processed using nf-core/rnaseq pipeline version 3.13.2^66^. The pipeline was executed with Nextflow version 23.10.0 on The Francis Crick Institute NEMO high performance computing cluster. Reads were aligned to the GRCh38 genome using Star. Gene expression quantification was performed using Salmon and quality control of raw and processed reads was performed with MultiQC. Differential gene expression analysis was performed using DESeq2 in RStudio running R v4.3.1.

### AlphaFold protein structural prediction

For protein modelling, AlphaFold 2 v2.3.1 was run on The Francis Crick Institute NEMO high performance computing cluster. The predicted aligned error (PAE) and predicted local distance difference test were analysed using UCSF ChimeraX and PyMOL to validate the model.

### Protein extraction and western blot

Cells were cultured and differentiated as described previously in a 6 well culture plate. Media was aspirated and the plate was placed directly in the −80°C freezer. For protein extraction, 500 µL Laemelli buffer solution (50 mM Tris-HCl pH 6.8, 20% glycerol, 2% SDS was added directly onto the culture dish and cells were scraped and transferred to an Eppendorf tube. Cells were than boiled at 95°C for 5 minute and placed on ice for 2-3 minutes. Samples were then centrifuged briefly. Protein quantification was carried out using BIORAD Protein Assay Kit (BIORAD, 5000002) as per the manufacturer’s protocol. Protein concentration was determined using BSA standards diluted in Laemelli buffer. Western blot was carried out using 40µg of protein for each sample. Samples were diluted with pre-made 4x loading buffer (BIORAD, 1610747), supplemented with DTT to a final concentration of 1 mM and boiled for 5 minutes at 95°C before briefly centrifuging and placing on ice. Samples were run on 4-20% Mini-PROTEAN TGX Precast protein gels (BIORAD, 4561094). Gels were run at 100 V for 10 minutes and 120 V thereafter for 2 hours in protein running buffer with 1% SDS. Samples were transferred to nitrocellulose membranes using iBlot2 system (Invitrogen, IB24001) for 6 minutes at 25 V according to manufacturer’s protocol. Membranes were blocked for 1 hour in 5% w/v milk diluted in Tris-buffered saline with tween 20 (TBST; 20mM Tris ph7.5, 150mM NaCl, 1% (v/v) tween 20). Lamin A/C primary antibody (SLS, SAB4200236-200UL) was used at a concentration of 1:500 in milk overnight at 4°C on a gentle rocker. The next day, membranes were washed 3 times for 5 minutes with TBST. Secondary antibody (Goat Anti-Mouse IgG (H + L)-HRP Conjugate (BIORAD, #1706516)) was diluted in 5% milk diluted in TBST for 1 hour at 4°C on a gentle rocker. Membranes were washed with TBST 3 times for 5 minutes. Bands were visualised using Amersham ECL western Blotting Detection Reagents (GE Healthcare, 28980926) according to the manufacturer’s instructions and imaged using BIORAD Chemidoc MP Imaging System.

### Immunostaining

Cells were fixed with 4% paraformaldehyde (PFA; Boster, AR1068) for 10 minutes at room temperature and washed with PBS. Cells were permeabilized in PBS with 0.2% Trition-X-100 (Sigma Aldrich, T8787-50mL) and 1% bovine serum albumin (BSA; Sigma Aldrich, A3294-50G) for 40 minutes at room temperature. Cells were blocked for 40 minutes at room temperature with permeabilization buffer supplemented with 10% goat serum (Sigma Aldrich, G9023-10mL). Primary antibodies were diluted in permeabilization buffer and incubated overnight at 4°C. The next day, primary antibodies were removed and cells washed 3 times for 5 minutes in PBS with 0.2% Triton-X-100. Secondary antibodies were diluted in the same solution and incubated for 1 hour at room temperature while blocked from light. Afterwards, cells were washed 3 times with PBS and stored at 4°C until imaged.

Engineered muscles were immunolabelled as previously described^57^. Briefly, muscles were fixed in 4% PFA while still attached to posts for 3 hours at 4°C, then washed with PBS before being removed from posts and placed into blocking buffer (0.05 M TBS (1x) pH7.4, 10% FBS, 1% BSA, 0.5% Triton-X-100) for 6 hours. Primary antibodies were incubated in blocking buffer without 10% FBS overnight on a shaker at 4°C. The next day, 6 washes were applied throughout the day with 1x TBS. Secondary antibodies were diluted in antibody buffer overnight on a shaker at 4°C. The next day, muscles were washed 6 times with 1x TBS before the muscles were mounted onto cavity slides (MERCK Life Science Ltd. UK, BR475505-50EA) with mounting medium (SIGMA Aldrich, F4680-25mL).

Primary antibodies used in this study were: rabbit anti NANOG, 1:100 (Abcam ab800892); mouse IgG2B anti OCT3/4, 1:100 (Santa Cruz Biotechnology sc5297); rabbit anti SOX2, 1:200 (Abcam ab92494); mouse IgG2B anti Lamin A/C, 1:250 (Invitrogen MA3100); rabbit anti Lamin B1, 1:100 (Abcam ab16048); mouse IgG1 anti MYH3, 1:9 (DSHB F1.652); rabbit anti NUP153, 1:100 (Abcam ab84872); mouse IgM anti Titin, 1:100 (DSHB 9D10). Secondary antibodies used were all AlexaFluor (Life Technologies 1:500). When immunostain 3D engineered muscles, all antibody dilutions were halved.

### Nascent RNA labelling

Nascent RNA was labelled using Click-iT^TM^ RNA AlexaFluor 594 Imaging Kit (Thermo Fisher Scientific, C10330). Briefly, cells were differentiated into myotubes as previously described on µ-Plate 24 Well ibiTreat (Ibidi, 82426). Cells were incubated with 1 mM EU for 1 hour prior to fixation for 15 minutes in 4% PFA. Cells were washed with PBS and permeabilised with 0.5% Triton-X for 1 hour. Click-IT reaction cocktail was added to the cells for 30 minutes before washing with Click-IT reaction rinse buffer. Immunostain for Lamin A/C and embryonic myosin heavy chain was then carried out as described previously.

### Microscopy

For standard fluorescence microscopy, a Nikon Ti2 inverted microscope was used with Teledyne Photometrics Prime BSI scientific CMOS camera and ASI motorised XY stage with Piezo Z, controlled with Micro-Manager v2.0 software. Imaging was carried out using either Plan Fluor 10x 0.30NA Ph1 or Nikon CFI S Plan Fluor ELWD 20XC lens with 1.5x zoom for final magnification of 30x for nuclear shape measurements. Spinning disc imaging for analysis of Lamin A/C and Lamin B1 mislocalisations and nucleoplasmic reticulum were conducted on a Nikon Ti2 with Yokogawa CSU-W1 SORA scanhead, driven with NIS-Elements using 40x lens and no additional zoom. The same microscope was used for imaging of 3D engineered muscles for the purpose of nuclear length measurements. For imaging of iPSC pluripotency markers in *LMNA* L302P iPSCs, a Leica DM16000B inverted microscope was used.

### Analysis of nuclear morphology and Lamin A/C localisation

Nuclear abnormalities were quantified by circling individual nuclei using the polygon tool in Fiji imaging software. For each nucleus the circularity (4π area/circumference2), major length, perimeter and area of the selected region was measured as described previously^67^. Nuclear abnormality classification was carried out in a blinded manner to prevent bias. Classes were calculated using the cell counter tool in Fiji imaging software. Classes are as follows: “Normal”, “Deformed”, “Blebbed”, “Elongated” or “String”. “Elongated” nuclei were classified as any nucleus above 30 µm which was validated empirically as being longer than myonuclei observed in control lines. Nuclear length measurements were conducted using the line segment tool in Fiji.

Analysis of lamin mislocalisations was carried out in a blinded manner. Lamin A/C mislocalisations were typically observed as either aggregations, patchy signal or absence of signal in particular areas (usually at the nuclear envelope). Lamin B1 mislocalisations were typically characterized by absence of signal in particular areas such as in locations of Lamin A/C aggregates. In control nuclei, Lamin A/C and Lamin B1 signal perfectly overlap each other and so any Lamin A/C or Lamin B1 signal that did not overlap with one another in *LMNA*-mutant myonuclei were classified as being mislocalised.

To assess presence of nucleoplasmic reticulum, spinning disc Z-stacks were visualized as 3D structures using Imaris imaging software. Nucleoplasmic reticulum was identified as co-stained Lamin A/C and Lamin B1 signal traversing the nuclear interior. Nuclear folds, characterised by a long stretch of intranuclear Lamin signal, were not classified as nucleoplasmic reticulum. The same image was also opened in Fiji imaging software and after visual examination in Imaris, myonuclei with and without nucleoplasmic reticulum were counted and quantified blindly. Myonuclear length in 3D artificial muscle constructs were analysed using the line segment tool in Fiji imaging software. Lines were drawn from pole to pole of the nuclei along their major axis.

### Statistical analyses

Statistics were performed in R (v.4.3.1) and GraphPad PRISM. Statistical comparisons and post-hoc tests used for pairwise comparisons are indicated within figure legends. For all statistical comparisons, a minimum of three biological replicates were performed.

## AUTHOR CONTRIBUTIONS

Conceptualization: FST; PSZ; DM. Data curation and analysis: DM; HBSS; LP; VML; LR; AA; SJ; CTYW; ACW; PSZ; FST. Funding acquisition and project coordination: FST. Investigation and methodology: DM; HBSS; LP; VML; LR; AA; SJ; CTYW; ACW. Project administration: FST. Resources: GB; PSZ, FST. Supervision: FST; PSZ. Writing and visualization: DM and FST, with PSZ and input from all co-authors.

## ACKNOWLEDGEMENTS

We are grateful to Dr J.M. Cuisset (Centre de Référence des maladies rares neuromusculaires Nord/Est/Ile-de-France, Lille, France), Cure CMD, FUJIFILM Cellular Dynamics, L-CMD patients and their families for providing cells. We thank members of the Tedesco lab for feedback and discussion, F. Torri, N. Khokhar, S. Maffioletti, M. Renshaw (Crick CALM STP), J. Campbell (Crick Bioinformatics STP) and the Crick Flow Cytometry and Genomics STPs for experimental support and Dr. R. Ben Yaou (Centre de Recherche en Myologie) for maintenance of the *LMNA* UMD database. This work was funded by: • Muscular Dystrophy UK (19GRO-PS48-0188-1) • The European Union (EU; ERC no. 759108 – HISTOID; Horizon Europe project no. 101080690 – MAGIC). Views and opinions expressed are however those of the authors only and do not necessarily reflect those of the EU or HADEA. Neither the EU nor HADEA can be held responsible for them. This work is funded by the UK Research and Innovation (UKRI) under the UK government’s Horizon Europe funding guarantee grant no. 10078461, 10080927 and 10079726; • LifeArc (CMCK-UK.FID119430465); • AFM-Téléthon (29063) • Biotechnology and Biological Sciences Research Council (BB/J014567/1; BB/M009513/1). FST also acknowledges support by Great Ormond Street Hospital Charity, Myotubular Trust, RYR1-Foundation, The Francis Crick Institute (which receives its funding from Cancer Research UK, the UK Medical Research Council and the Wellcome Trust) and the National Institute for Health and Care Research (NIHR; the views expressed are those of the authors and not necessarily those of the National Health Service, the NIHR, or the Department of Health). Graphical abstract created with BioRender.com. For the purpose of Open Access, the author has applied a CC BY public copyright licence to any Author Accepted Manuscript version arising from this submission.

