## Supplemental Information for "Advanced human iPSC-based modelling of *LMNA*-related congenital muscular dystrophy enables development of targeted genetic therapies for muscle laminopathies"

### Supplementary figures

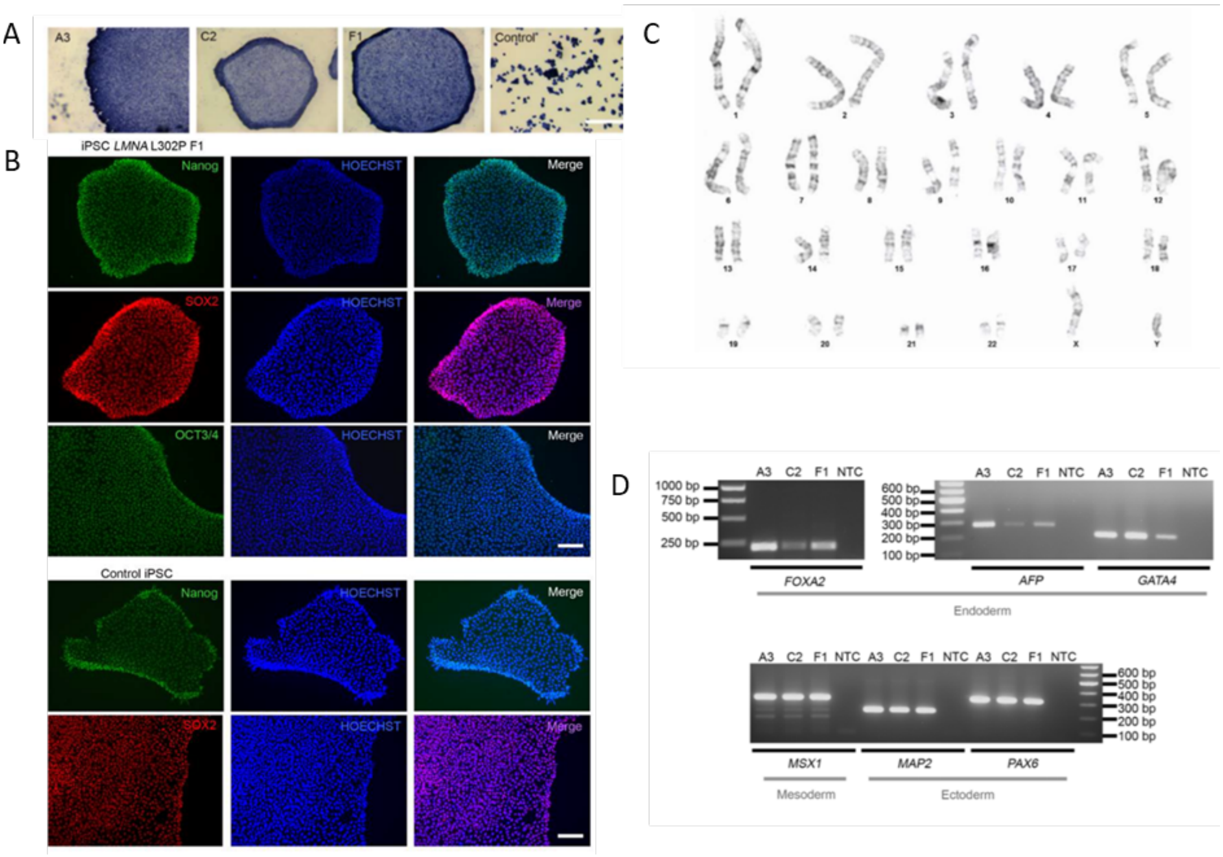


**Supplementary figure 1. Characterisation of *LMNA*$^{L302P}$ iPSCs.**

**(A)** Alkaline phosphatase staining of *LMNA* L302P iPSCs. A3, C2 and F1 represent individual clones following reprogramming. HeLa cells included as positive control. Scale bar 500µm. **(B)** Representative immunofluorescence for pluripotency markers SOX2, OCT3/4 and NANOG in reprogrammed *LMNA* p.L302P iPSCs F1 clone, which was used in this work. Scale bar 100 µm. **(C)** Karyotype analysis of *LMNA* L302P F1 clone. Normal male chromosome complements and banding pattern visible. **(D)** Reverse transcription PCR for the markers of the three germ lineages endoderm, mesoderm and ectoderm following spontaneous differentiation of iPSCs in DMEM 20 %. Three endoderm (FOXA2, AFP, GATA4), one mesoderm (MSX1) and two endoderm markers (MAP2 and PAX6) amplify and produce PCR products of the expected size in all three of the differentiated *LMNA* p.L302P iPSC lines, demonstrating their three-lineage pluripotency. FOXA2 = 216 bp cDNA and 1253 bp genomic DNA, AFP = 281 bp cDNA and 1939 bp genomic DNA, GATA4 = 219 bp cDNA and 1428 bp genomic DNA, MSX1 = 307 bp cDNA and 2639 bp genomic DNA, MAP2 = 212 bp cDNA and 431 bp genomic DNA, PAX6 = 317 bp cDNA and 644 bp genomic DNA. Assay was run once per iPSC clone. A no template control was run with the reaction to control for genomic/PCR product contamination, which produced no PCR product. NTC = no template control.

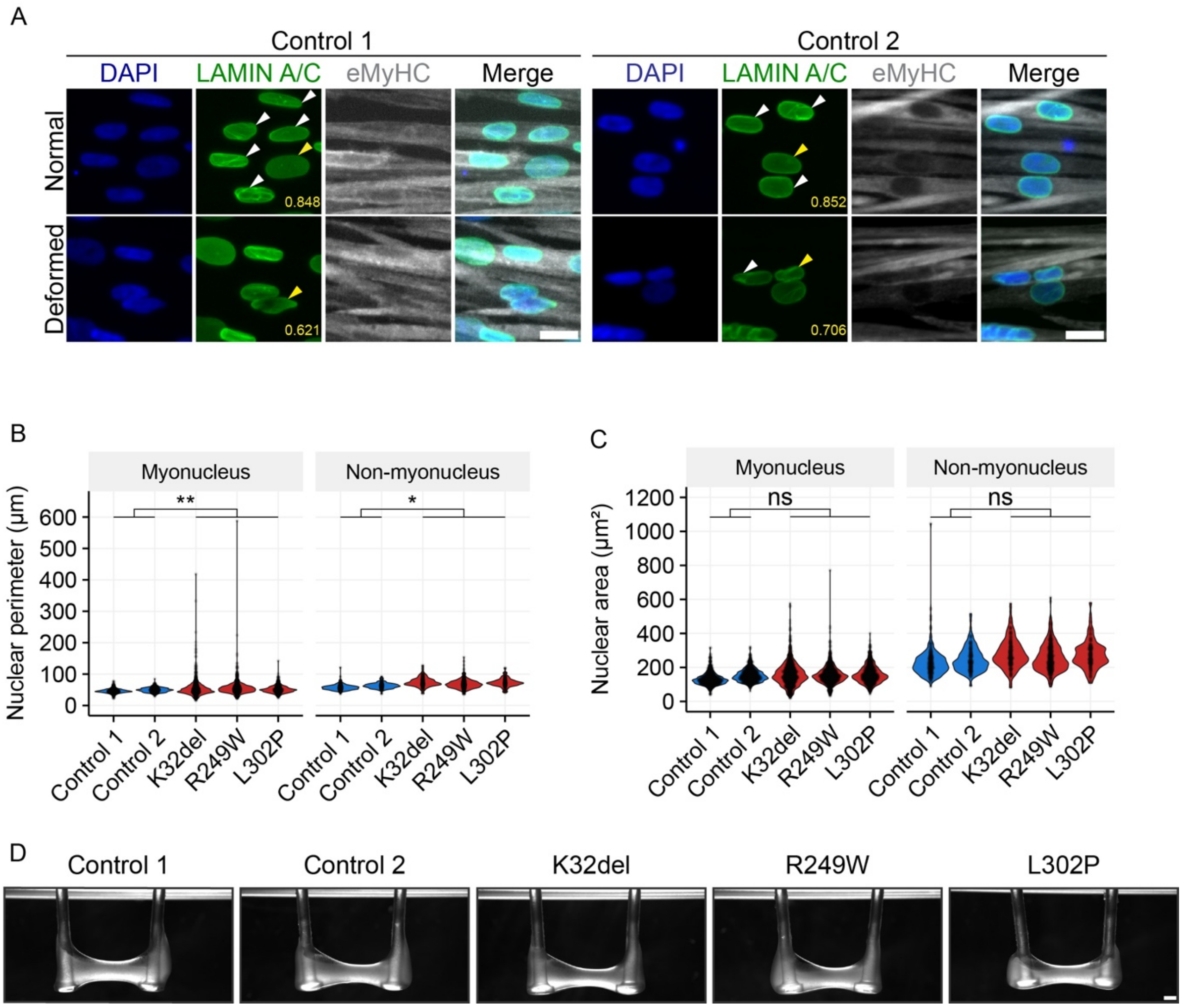


**Supplementary figure 2. Additional characterisation of skeletal myogenic progenitors derived from *LMNA*-mutant iPSCs.**

**(A)** Immunostaining of control cell lines for Lamin A/C and embryonic myosin heavy chain (eMyHC) showing representative images of five nuclear morphology classifications are displayed: Normal, Deformed, Blebbed, Elongated, and String. Arrows indicate nuclei belonging to shape classification. Yellow arrows indicate nucleus associated with nuclear circularity measurement in lower right corner. Scale bar 20 µm. **(B, C)** Graphs showing quantification of (A) nuclear perimeter and (B) area. One-way ANOVA with Bonferroni correction for pairwise comparisons. Data presented as scatter plots showing measurements of all nuclei across repeats with violin plots showing distribution of measurements. Three independent passages were analysed per cell line (N=3). 205-622 myotube nuclei and 50-174 non-myotube nuclei were analysed per cell line, per repeat. ns p≥0.05; * p<0.05; ** p<0.01. **(D)** Stereoscope images of 3D engineered muscles. Scale bar 100 µm.

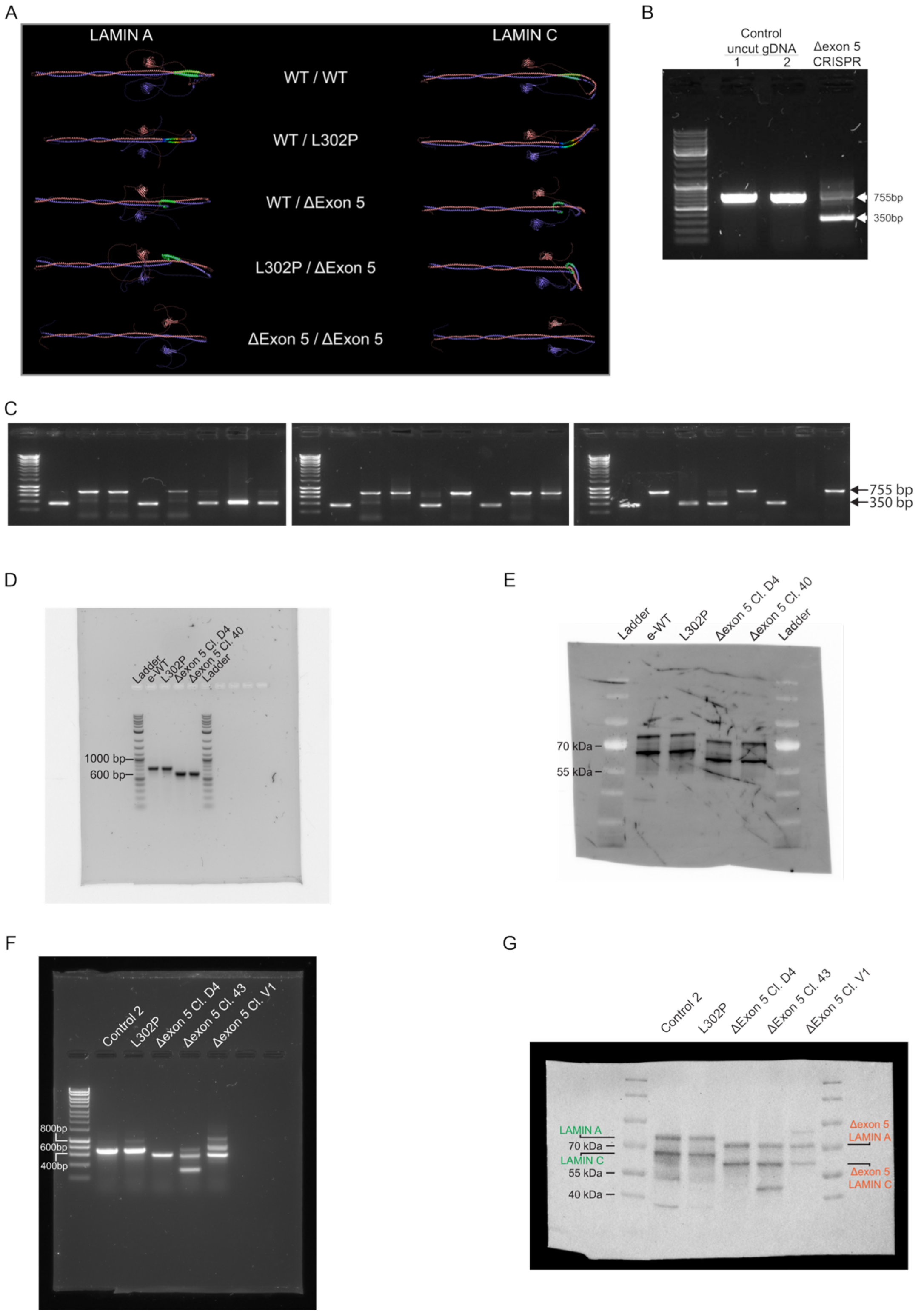


**Supplementary figure 3. Additional molecular characterisation of CRISPR edited *LMNA*-mutant cells.**

**(A)** AlphaFold structural predictions of WT, L302P and Δexon 5 Lamin A/C dimerization. **(B)** PCR of *LMNA* gene showing control uncut gDNA and Δexon 5, CRISPR-treated, pooled gDNA. 755 bp shows uncut sequence. 350 bp shows exon-removed gDNA. **(C)** PCR of *LMNA* gene in clonally expanded Δexon 5 iPSCs. **(D)** Full agarose gel of PCR of cDNA from CRISPR-corrected (e-WT), *LMNA* L302P, and

Δexon 5 (clones D4 and 40) myogenic cell. **(E)** Full membrane of western blot for Lamin A (upper band) and Lamin C (lower band) from CRISPR-corrected (e-WT), *LMNA* L302P, and Δexon 5 (clones D4 and 40) myogenic cells.**(F)** PCR of cDNA from Δexon 5 clones D4, 43 and V1 myogenic cells, showing additional undesired *LMNA* bands as a result of recombination events following CRISPR in clones 43 and V1. **(G)** Western blot for Lamin A/C from Δexon 5 clones D4, 43 and V1 myogenic cells showing additional undesired protein products as a result of recombination events following CRISPR treatment.

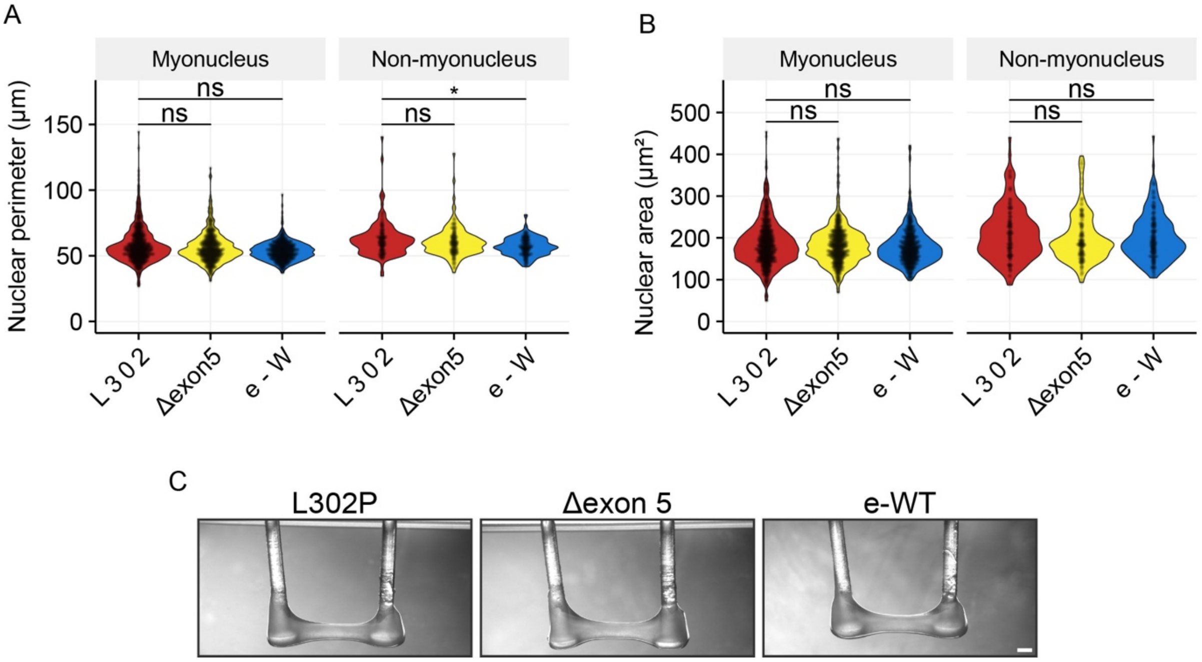


**Supplementary figure 4. Additional characterisation of skeletal myogenic progenitors derived from CRISPR-edited *LMNA*-mutant iPSCs.**

**(A, B)** Graphs showing quantification of (A) nuclear perimeter, (B) nuclear area. Beta regression with Bonferroni correction for pairwise comparisons. Data presented as scatter plots showing measurements of all nuclei across repeats with violin plots showing distribution of measurements. Three independent passages were analysed per cell line (N=3). 181 - 301 myotube nuclei and 32 - 108 non-myotube nuclei were analysed per cell line, per repeat. ns p≥0.05; * p<0.05. **(C)** Stereoscope images of 3D engineered muscles. Scale bar 100 µm.

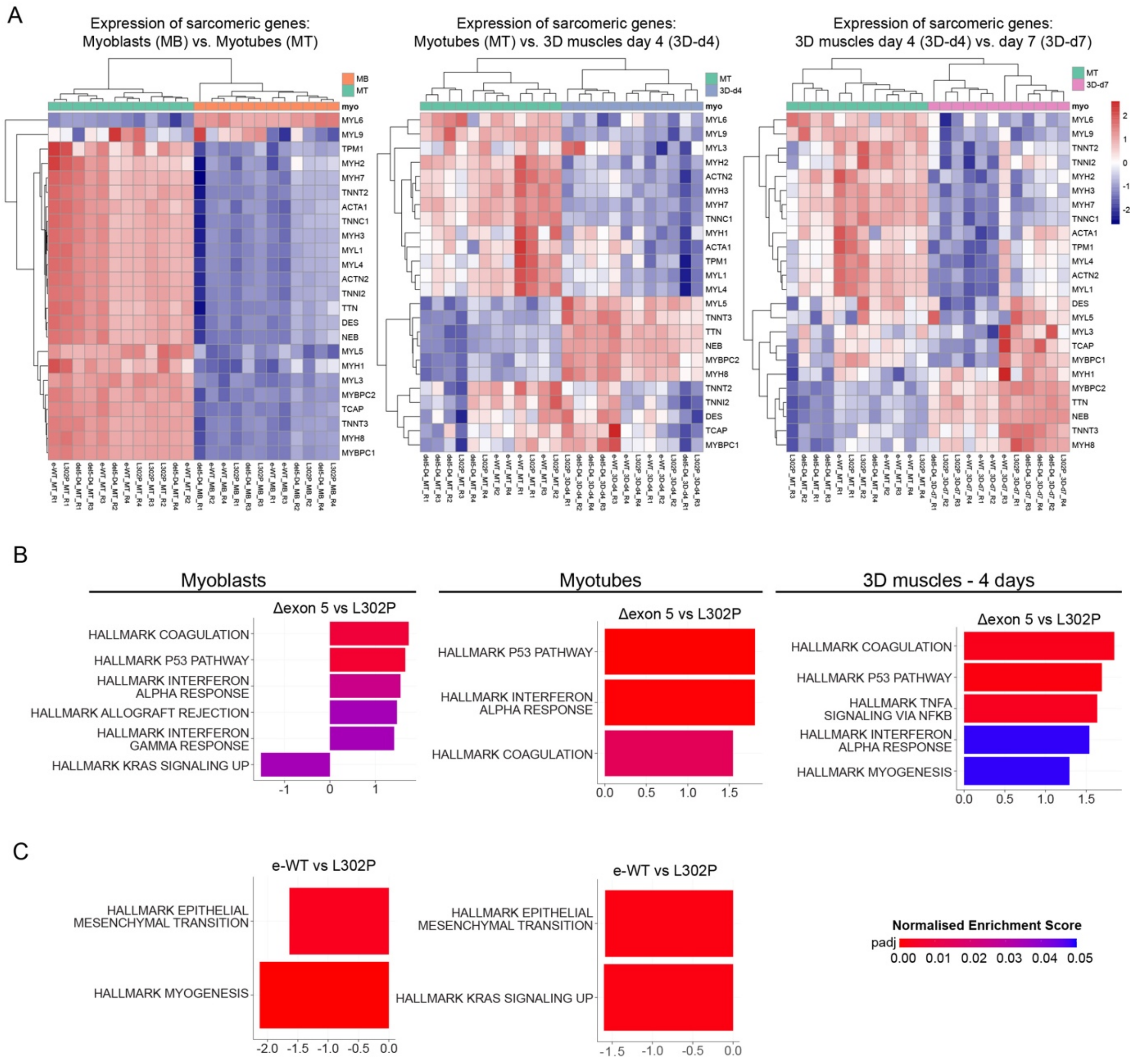

**Supplementary figure 5. Additional transcriptomic characterisation of skeletal myogenic progenitors derived from CRISPR-edited *LMNA*-mutant iPSCs: maturation and gene set enrichment analysis.**

**(A)** Heatmaps showing normalised expression of sarcomeric genes in myoblasts (MB) and 3D engineered muscles differentiated for 4 days (3D-d4) and 7 days (3D-d7) respectively, using 2D myotubes (MT) as a reference. **(B, C)** GSEA of **(B)** ∆exon 5 vs L302P and (**C)** e-WT vs L302P in myoblasts, myotubes and 3D engineered muscles at day 4 of differentiation. Indicated hallmark gene sets used in Figure 6D were analysed. All significantly enriched gene sets (padj < 0.05) in each comparison are shown. Note, no enriched pathways in e-WT vs L302P in 3D muscles at 4 days of differentiation.

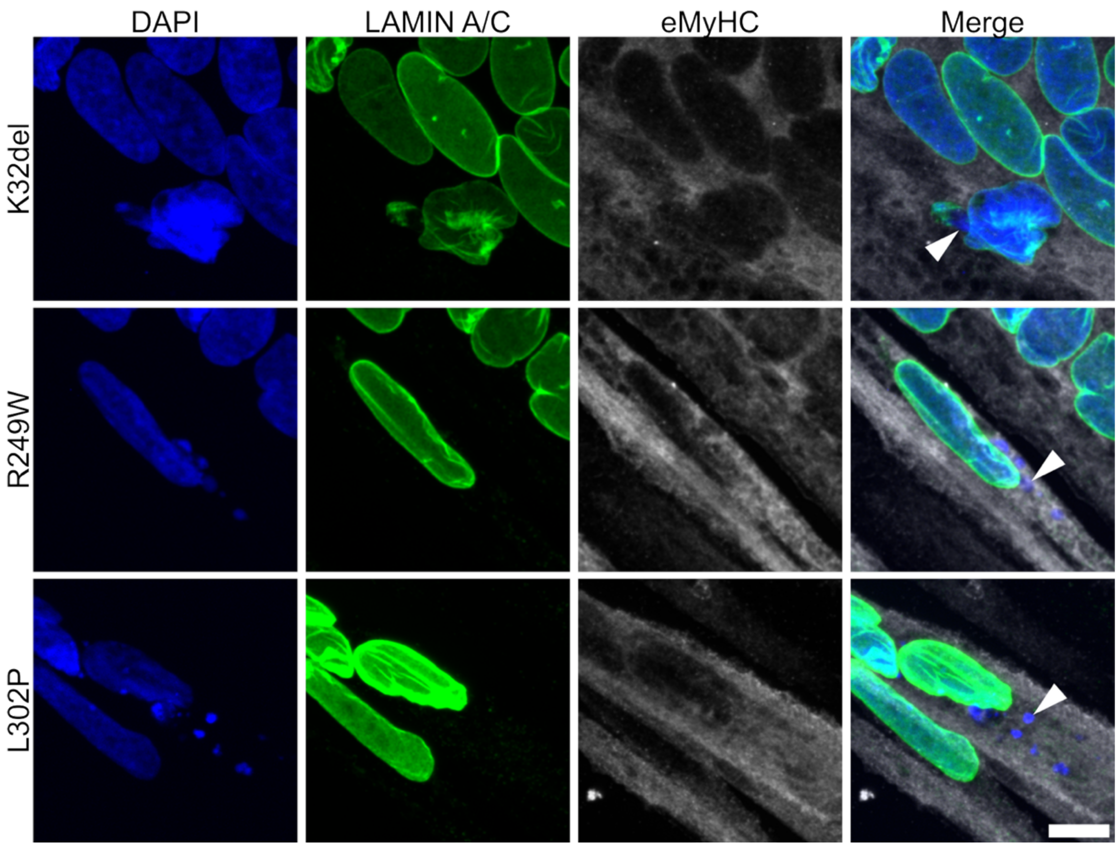


**Supplementary figure 6. Cytoplasmic DNA in *LMNA*-mutant iPSC-derived myotubes.**

Immunostaining of *LMNA*-mutant myotubes for Lamin A/C and embryonic myosin heavy chain, counterstained with DAPI. White arrows show DAPI signal leaking from myonuclei. Scale bar 10 µm.

**Supplementary tables**

| Homology Directed Repair | | |
|---|---|---|
| ***LMNA* mutation** | **sgRNA** | **ssODN repair template** |
| L302P (c.905T>C) | CTTCTGGAGCTGGCTG<br>AGCT | GCAACCTGGTGGGGGC<br>TGCCCACGAGGAGCTG<br>CAGCAGTCGCGCATCC<br>GCATCGACAGCCTCTCT<br>GCACAGCTCAGCCAGC<br>TCCAGAAGCAGGTGATA<br>CCCCACCTCACCCCTCT<br>CTCCAG |

**Supplementary table 1.** sgRNA and ssODN homology directed repair template used for correction of *LMNA* L302P mutation. Green highlight nucleotide indicates correction. Cyan highlighted nucleotide indicates silent PAM blocking mutation.

| ***LMNA* L302P permanent exon skipping** | | |
|---|---|---|
| **sgRNA Name** | **Location** | **Sequence** |
| sgRNA 1 | Intron IV | CCAGAAGGCATAGCCCAGCG |
| sgRNA 2 | Intron V | GACCCTTCTCTGTGGTTGTG |

**Supplementary table 2.** sgRNAs used for generation of Δexon 5 line.

<table>
<tr><th colspan="3">Primers used</th></tr>
<tr><td>Purpose</td><td>Forward</td><td>Reverse</td></tr>
<tr><td>K32del PCR</td><td>AGAGGAGGACCTATTAGAGC</td><td>CCCACCATTCCTTATATCCTC</td></tr>
<tr><td>R249W PCR</td><td colspan="2" rowspan="3">GGTTTCTGTGTCCTTCCTCC<br>CCTGAGAAGTGAAGGTGAGG</td></tr>
<tr><td>L302P PCR</td></tr>
<tr><td>Δexon 5 gDNA PCR</td></tr>
<tr><td>Δexon 5 cDNA PCR</td><td>CCTGATAGCTGCTCAGGCTC</td><td>GGCCAGCTTGATGTCCAGAA</td></tr>
</table>

**Supplementary table 3.** Primers used in this study.
